# Unravelling Induced Resistance in Strawberry: Distinct Metabolomic Signatures Define Cultivar-Specific Resistance to *Botrytis cinerea*

**DOI:** 10.1101/2025.07.29.667391

**Authors:** Chiara Murena, Victoria Pastor, Tânia R. Fernandes, Susana M.P. Carvalho, Estrella Luna

## Abstract

*Botrytis cinerea* is a major pathogen in strawberry, and sustainable alternatives to fungicides are needed to manage this disease. Induced resistance (IR) through chemical elicitors represents a promising strategy, but the effectiveness of such compounds remains poorly understood in commercial strawberry (*Fragaria × ananassa*) cultivars. In this study, we evaluated the efficacy of repeated applications of five elicitors (*i.e*., β-aminobutyric acid (BABA), (R)-β-homoserine (RBH), indole-3-carboxylic acid (I3CA), jasmonic acid (JA), and salicylic acid (SA)) in three strawberry cultivars (Rowena, Soraya, and Durban). BABA and RBH significantly reduced B. cinerea lesion sizes in Rowena and Soraya, while Durban showed no induced resistance to the elicitors. Untargeted metabolomic profiling of Rowena and Soraya revealed cultivar-specific responses to elicitor treatment and infection, with distinct patterns of metabolite accumulation under both mock- and *B. cinerea*-inoculated conditions. RBH in Rowena and BABA in Soraya induced the most extensive priming-associated metabolic reprogramming, including enrichment of amino acid, nucleotide, and secondary metabolite pathways such as flavonoids and phenylpropanoids. Significantly, none of the elicitors negatively affected plant growth, flowering, or fruit set. These results demonstrate that the effectiveness and mechanism of IR in strawberries depend on both the elicitor and the cultivar, providing new insights into the metabolomic basis of priming with implications for sustainable disease management in strawberry cultivation.

## Introduction

The necrotrophic fungus *Botrytis cinerea*, the causal agent of grey mould disease, is among the most economically damaging pathogens affecting strawberry plants (*Fragaria × ananassa* Duch.; Family: *Rosaceae*) (Petrasch et al., 2019). Strawberries are a significant global commodity, with an annual production of approximately 10 million tons worldwide (FAO, 2023). However, this productivity is severely threatened because of the crop’s high susceptibility to *B. cinerea* (Bestfleisch et al., 2015). Grey mould manifests as necrotic lesions that rapidly develop into water-soaked areas, where the fungus spreads, producing dense mycelial growth and spores, ultimately leading to the collapse of the affected organs (Williamson et al., 2007). It is estimated that over half of the strawberry yield can be lost at postharvest if plants are not treated with fungicides before the harvest (Hassan et al., 2021, Petrasch et al., 2019). Despite the emergence of *B. cinerea*-resistant strains and the potential adverse effects on human health and the environment, chemical fungicides remain widely used due to their high efficacy (Chen et al., 2016). Consequently, it is crucial to develop new sustainable chemical alternatives to reduce *B. cinerea* incidence in strawberry. A deeper understanding of strawberry defence responses to *B. cinerea* is therefore essential. One promising strategy for sustainable crop protection is harnessing the plant immune responses using naturally derived compounds to stimulate Induced Resistance (IR). When exposed to local and temporary stress *stimuli*, plants can develop an IR phenotype, characterized by enhanced defence capacity and reduced susceptibility to future challenges (De Kesel et al., 2021). This response may involve direct activation of defences, which can impose severe plant fitness costs due to the constant expression of defence mechanisms, or a more energy-efficient process known as priming, where plants, after the perception of a first *stimulus*, are sensitised to respond more rapidly and strongly to subsequent attacks (He et al., 2022, Mauch-Mani et al., 2017, van Hulten et al., 2006). Priming is particularly attractive as it offers the benefits of enhanced defences with minimal fitness costs (Martinez-Medina et al., 2016). It can be long-lasting, creating a durable immunological memory that is maintained throughout the plant’s life cycle and even transmitted across generations, being particularly relevant for vegetatively propagated crops (Catoni et al., 2022), such as strawberry. Several exogenous chemical agents, known as elicitors, can trigger IR and prime the plant’s defences. For instance, the application of the phytohormones jasmonic acid (JA) and salicylic acid (SA) can prime plant defences (Pastor et al., 2013). Typically, JA mediates defence against necrotrophic pathogens, while SA is more effective against biotrophs (Ghozlan et al., 2020, Liao et al., 2022). JA has successfully primed tomato defences against *B. cinerea* when applied to seeds or seedlings, without causing any growth reduction (Luna, 2016, Worrall et al., 2012). On the contrary, the role of SA in defence against *B. cinerea* is unclear. While beneficial effects were observed following the application of SA in tomato (Li and Zou, 2017), pepper (Mekawi et al., 2019), and strawberry (Babalar et al., 2007), enhanced susceptibility was reported (El Oirdi et al., 2011, Fugate et al., 2013, Ha et al., 2021, Khanam et al., 2005). The contradictory findings, together with the limited investigation of JA and SA as preharvest treatments, highlight the necessity for specific case studies within the *B. cinerea*-strawberry pathosystem (Koo et al., 2020). The non-protein amino acid β-aminobutyric acid (BABA) has emerged as a potent chemical priming agent. Once considered a xenobiotic, BABA has recently been identified as a natural plant metabolite involved in stress signalling (Thevenet et al., 2017). BABA induces broad-spectrum resistance by priming multiple signalling pathways, including both SA-dependent (Zimmerli et al., 2001) and SA-independent mechanisms (Ton and Mauch-Mani, 2004). In *Arabidopsis thaliana*, its action is mediated by binding the active (R)-enantiomer of BABA to aspartyl-tRNA synthetase (AspRS), activating defence-related pathways (Luna et al., 2014a). BABA has shown efficacy in several crops, including the model plant *A. thaliana* (Koen et al., 2014, van Hulten et al., 2006, Zimmerli et al., 2001) and tomato (Koen et al., 2014, Luna et al., 2016, Luna et al., 2020, van Hulten et al., 2006, Wilkinson et al., 2017, Zimmerli et al., 2001) as well as in postharvest protection of grapevine (Csikász-Krizsics, 2013), onions (Polyakovskii et al., 2008), and strawberries (Wang et al., 2016). Its effect on the strawberry plant is highly variable, from enhanced resistance to increased susceptibility, depending on application method, BABA concentrations, and plant developmental stage (Badmi et al., 2022, Badmi et al., 2019).

Despite its potential, BABA application for crop protection has been hampered by its growth-inhibiting properties, which result from the inhibitory binding of BABA to AspRS enzymes, thereby competing with Asp and preventing its binding (Luna et al., 2014a). However, while some studies reported remarkable effects on growth and yield (Badmi et al., 2019, Buswell et al., 2018, van Hulten et al., 2006), others report just a transient growth reduction (Luna et al., 2014b, Wilkinson et al., 2017) or no effects (Badmi et al., 2019), highlighting the importance of species-specific evaluation. To overcome these drawbacks, BABA analogues, such as (R)-β-homoserine (RBH), have been investigated against *B. cinerea*. RBH retains IR-inducing properties without causing growth suppression and has been shown to prime JA defences in tomato and to provide long-lasting protection in *Fragaria vesca* plants (Badmi et al., 2019, Buswell et al., 2018). Another candidate priming agent is indole-3-carboxylic acid (I3CA), a metabolite that has been found to accumulate in BABA-primed *A. thaliana* upon infection with *Plectosphaerella cucumerina* (Gamir et al., 2012, Gamir et al., 2014). When exogenously applied, I3CA was effective in inducing resistance against *P. cucumerina* in *A. thaliana*, but the I3CA-induced resistance appeared to be independent of the SA and JA pathways (Gamir et al., 2014). Moreover, no study has been conducted to assess its effects on plant growth. Therefore, further studies are needed to understand the mechanisms underlying I3CA-induced resistance against necrotrophic pathogens and evaluate its suitability for crop protection. BABA, RBH, and I3CA showed no direct antifungal activity, supporting the idea that their efficacy is plant-mediated rather than due to directly killing the pathogen (Badmi et al., 2019, Buswell et al., 2018, Gamir et al., 2014). However, their efficacy in strawberry, as well as the underlying mechanisms, remains underexplored. Priming induced by elicitor application presents a sustainable and promising strategy for protecting crops from *B. cinerea*. Since priming involves complex biochemical changes, metabolomic approaches provide a powerful tool for unravelling the complex mechanisms behind defence priming. Identifying key metabolic pathways and markers associated with IR could facilitate the development of more sustainable chemical strategies. To date, a comprehensive metabolomic analysis to unravel IR mechanisms primed by SA, JA, BABA, RBH, and I3CA in strawberry against *B. cinerea* is lacking.

No fully resistant genotypes have been identified, and the genetic and biochemical factors contributing to resistance against *B. cinerea* remain largely unknown (Bestfleisch et al., 2015). For instance, the resistance of strawberry against *B. cinerea* is described as quantitative disease resistance (QDR), which leads to partial resistance, and its success may be highly affected by external factors, such as environment, plant species and variety, and plant developmental stages. On the other hand, the high pathogenicity of *B. cinerea* in strawberry arises from the ability to deploy a rich toolbox of nonspecific pathogenicity factors for which no known resistance (R) genes in strawberry are available (Bi et al., 2023). Moreover, non-genetic variables in field studies (e.g., row density, climatic conditions) have been considered as the primary cause of slight differential susceptibility (Rhainds et al., 2002).

Thus, this study investigates: (1) the efficacy of BABA, RBH, I3CA, JA, and SA in inducing resistance to *B. cinerea* in three commercial strawberry cultivars (Durban, Rowena, and Soraya); (2) the mechanisms of IR through untargeted metabolomics; (3) the metabolic pathways and key metabolites involved in priming; and (4) the potential growth reduction and fitness costs of elicitor treatments. Our results aim to provide novel insights into the defence responses of strawberries to *B. cinerea* and the mechanisms of defence priming agents, thereby developing new sustainable crop protection strategies.

## Materials and methods

### Plant Material and Growth Conditions

Seeds of three *Fragaria × ananassa* cultivars, Durban, Rowena, and Soraya, were obtained from seeds (ABZ seeds, UK, kindly provided by Saturn Bioponics https://saturnbioponics.com/). Seeds were germinated on wetted filter paper in sealed Petri dishes under long-day conditions (16 h light/8 h dark, 25°C/20°C, 100% relative humidity) in a growth chamber. After two weeks, uniformly germinated seedlings were transplanted into alveolar trays (80 mL volume per cell) filled with Levington M3 compost. Plants were grown under the same photoperiod, temperatures, and light intensity until they reached 18 weeks of age. Water and nutrient supply were adjusted according to plant phenological stages using the commercial fertigation protocols of Vitax Organic Strawberry Fertiliser (Catalogue number 5LSF1).

### Pathogen Culture and Inoculum Preparation

*Botrytis cinerea* (isolate BcI16) was cultured on potato dextrose agar (PDA) plates. A single agar plug from a sporulated colony was transferred to fresh PDA and incubated in the dark at room temperature for four weeks as previously described (Stevens et al., 2025). Spores were collected in sterile water containing 0.01% Tween-20, filtered through Miracloth® (EDM Millipore, Burlington, MA, USA), and pelleted by centrifugation (10 min at 4,000 rpm). The final spore concentration of the inoculum was adjusted to 5 × 10⁵ spores/mL in half-strength potato dextrose broth (PDB).

### Elicitor Treatments

Five elicitors were tested: β-aminobutyric acid (BABA), (R)-β-homoserine (RBH), indole-3-carboxylic acid (I3CA), jasmonic acid (JA), and salicylic acid (SA). Compounds were obtained from Sigma-Aldrich and prepared freshly prior to application. BABA and RBH were applied at 0.5 mM, I3CA at 0.150 µM, JA at 0.1 mM, and SA at 1 mM final concentrations. BABA, RBH, and I3CA were applied via soil drenching by injecting a 10x concentrated solution at 10% of the tray’s alveolus volume (80 mL). BABA and RBH were dissolved in sterile distilled water (SDW). I3CA was dissolved in ethanol and then diluted to the selected concentration. The final ethanol concentration in the solution was 0.075%, and therefore, all soil drench solutions were supplemented with ethanol to achieve this concentration. JA and SA were applied as foliar sprays until runoff on both adaxial and abaxial leaf surfaces. SA was dissolved in SDW. Stock solutions of JA were prepared according to Luna et al. (2016) by dissolving 250 mg in 2 ml of ethanol, then diluted in distilled water to a final stock concentration of 10 mM, and stored at -20 °C. Before use, the 10 mM stock solution was thawed and diluted to the final concentration in the spraying solution, resulting in a final ethanol concentration of 0.042%. Therefore, all other spraying solutions were supplemented with 0.042% ethanol. All foliar spray solutions contained 0.01% Silwet to improve adherence to leaf surfaces. Solvent solutions only (without chemicals) were used to treat control plants (*i.e*., Water treatment), and the soil-drench solution and spraying solution were applied to each treatment to equalise the amount of solvent in the soil or on the leaves. All treatments were applied four times at 2-week intervals on plants aged 6, 8, 10, and 12 weeks (Figure 1A).

**Figure 1.**
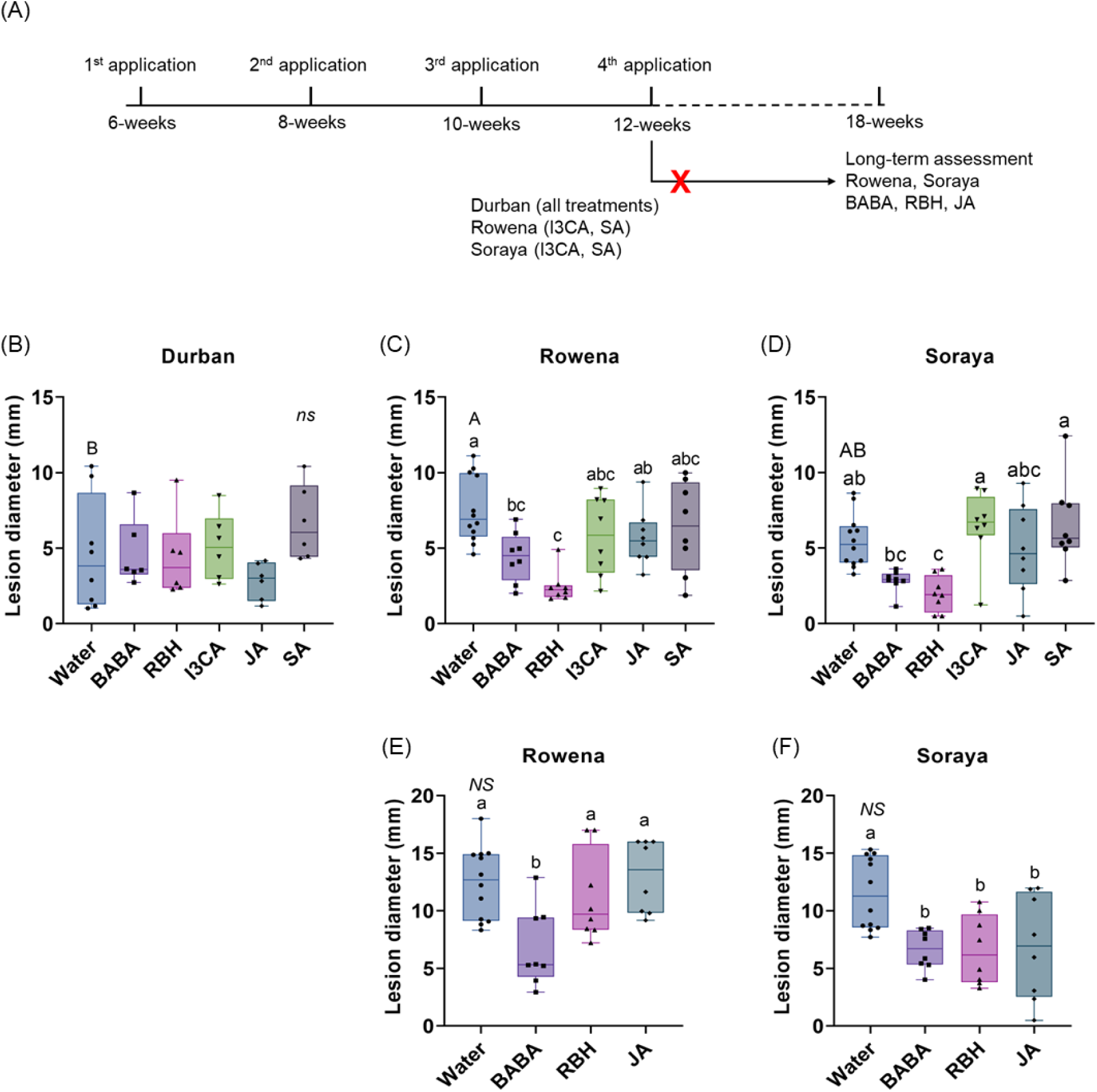
Induced resistance to *B. cinerea* in strawberry cultivars treated with chemical elicitors. (A) Experimental timeline showing four elicitor applications at 6, 8, 10, 12, and 18 weeks of plant age and subsequent leaf inoculations with B. cinerea for disease assessment. The red cross indicates the treatments that were excluded from the long-term assessment. Lesion diameter in Durban (B), Rowena (C), and Soraya (D) after four elicitor applications. Lesion diameter six weeks after the final elicitor application (18-week-old plants) in Rowena (E) and Soraya (F). Boxplots represent lesion diameter (mm) with the median line, interquartile range (boxes), and whiskers extending to the minimum and maximum values. Each point represents biological replicates (individual plants) (n = 8-12 per treatment). Capital letters indicate statistical differences between the Water treatments of each variety. Lowercase letters indicate statistically significant differences between elicitors within each variety (One-way ANOVA followed by Tukeýs post hoc test or Welch’s ANOVA followed by Dunnett’s T3; p < 0.05; n = 8–12).

### Leaf infection assay with *B. cinerea*

Detached leaf assays were conducted five days after each treatment application. One fully expanded leaf was selected for infection in each plant. Detached leaves (n = 8-12) were infected by applying 5 μL of the drop inoculum solution on either side of the main vein (*i.e*., 6 drops/leaf). Mock leaves were infected with ½ strength PDB only. Inoculated leaves were incubated at 20°C in the dark at 100% RH. Disease incidence was measured as necrotic lesion diameter (mm) five days post-inoculation using an electronic calliper (0.1 mm resolution).

### Growth and Reproductive Performance Assessment

Relative growth rate (RGR) for leaf area was calculated between the time points one week before the fourth elicitor application (t1; week 11 of the experiment) and two weeks after the same application (t2; week 14 of the experiment). The following formula was used: RGR = [ln(leaf area1₂) – ln(leaf area1₁)] / (t₂ – t₁). Leaf area was measured with ImageJ Software (Version 1.54j). Flowering and fruit production were recorded on 18-week-old plants (six weeks after the fourth application) by counting the number of flowers and fruit per plant.

### Statistical Analysis

Data normality was assessed using the Shapiro-Wilk test (p ≥ 0.05). Homogeneity of variance was evaluated with Levene’s test. For normally distributed data with equal variances, one-way ANOVA followed by Tukey’s post hoc test was used. When variances were unequal, Welch’s ANOVA followed by Dunnett’s T3 test was applied. Analyses were performed in GraphPad Prism 9 (GraphPad Software, San Diego, CA, USA).

### Metabolomic analysis

#### Sample collection for metabolomic analysis

For metabolomic analysis, leaves were collected from individual plants (n = 8-12) of Rowena and Soraya, following treatment with Water, BABA, RBH, and JA 5 days after the final application. Infected and mock-infected leaves were collected 24 hours post-infection (hpi), snap-frozen in liquid nitrogen, ground in liquid nitrogen, and lyophilized before metabolite extraction. For each condition, three biological replicates were used. Each biological replicate consisted of a pool of one leaf per plant (i.e., leaves from four plants for the water treatments and two to three plants for the chemical treatments).

#### Metabolite extraction and LC-QTOF analysis

Three biological replicates and three techniques were used for the analysis (n = 6). The metabolites were extracted using a mixture of MeOH and H_2_O (30:70) supplemented with 0.01% HCOOH. Then, 1 mL of 30% MeOH was added to 300 mg of powdered freeze-dried tissue, and the samples were incubated on ice for 30 min. After shaking for 15 min and centrifugation at 15.000 rpm for 15 min at 4°C, the supernatant was filtered using a 0.2 µm cellulose filter. An aliquot (5 µL) of each sample was injected into the hybrid tandem UPLC-QTOF (Synapt, Waters) mass spectrometer in the positive (ESI +) and negative (ESI −) ion modes for electrospray ionization. Samples elution was performed through a reversed-phase C-18 column (Kinetex EVO C18 Core-Shell, 2.6 µm particle size, Phenomenex) with a gradient of MeOH and H_2_O supplemented with 0.01% HCOOH. Raw data can be found in Metabolights (ID: data available upon acceptance).

Raw data were transformed into. mzML format using the MSConvert tool (ProteoWizard). Data were processed using R software version 4.4.1, where ESI+ and ESI-signals were analysed separately. Signal corrections were obtained using the Centwave algorithm for R. The amount of each compound was determined from the normalized peak area relative to the dry weight of each sample.

#### Global data visualization and statistical analysis

Global visualization of the combined ESI+ and ESI-data was performed using Principal Component Analysis (PCA) and a heatmap in Metaboanalyst 6.0, applying filtering through the interquartile range (<40%), normalization by median, followed by cube root transformation and Pareto scaling for combined positive and negative ionization mode data. For PCA, statistical significance between the treatment groups was evaluated using PERMANOVA (p ≤ 0.05).

Heatmaps were obtained using hierarchical clustering, Euclidean distance measurement, and Ward’s clustering method, and metabolite expression was represented as group averages. A log_2_ fold change measure was used to determine up- or down-expression levels of metabolites. Venn diagrams were drawn using Venny v2.1.0, and the significant putative metabolites associated with each treatment and infection condition were determined using a two-sample t-test (FDR, p ≤ 0.05) between the elicitor and Water treatments under both mock and B. cinerea infection.

#### Pathway enrichment analysis of priming metabolites

Priming metabolites were isolated from Venn diagrams by comparing the elicitor vs. Water upon mock infection and the elicitor vs. Water upon pathogen infection and selecting significant metabolites solely associated with infection. These metabolites were putatively identified through several metabolite libraries, one available in the KEGG database for *Fragaria vesca* and one internal library kindly provided by Dr. Pastor’s group. The priming metabolites specific to each compound underwent enrichment pathway analysis using the MarVis-Suite software. Adduct and isotope correction, followed by merging of the positive and negative ionization mode data, was performed using the MarVis-Filter, while the pathways were obtained from the MarVis-Pathway. Entry-based enrichment analysis calculated p-values based on a hypergeometric distribution, which were then adjusted using FDR (Benjamini-Hochberg) correction. Pathways with p-values ≤0.05 were considered significantly enriched. The putatively annotated Metabolites of enriched pathways were further identified at a confidence level (MS1 MS2) in MassLynx through ChromaLynx. When fragmentation spectra were not available in the selected libraries, fragmentation spectra from online databases (PubChem, HMDB) were considered, with similar LC-MS/MS experimental conditions and ionization modes.

## Results

### Efficacy of elicitors in inducing resistance to *B. cinerea*

To evaluate the ability of five chemical elicitors (*i.e*., BABA, RBH, I3CA, JA, and SA) to induce resistance against *B. cinerea*, four applications were carried out at two-week intervals on strawberry cultivars Durban, Rowena, and Soraya at 6, 8, 10, and 12 weeks of age (Figure 1A). A leaf infection assay was carried out five days after the application of the elicitors. Following inoculation, disease incidence was measured as lesion diameter (mm) five days post-infection.

Before elicitor analysis, basal resistance levels among the three cultivars were first assessed for Water-treated controls. No significant differences were observed after the first application (Supplementary Figure 1A). However, after the second application, Rowena exhibited higher susceptibility than Soraya, while Durban showed an intermediate response (Supplementary Figure 1B). These differences were not observed after the third application (Supplementary Figure 1C). However, after four applications, Rowena exhibited significantly higher susceptibility compared to Durban, indicating the lowest basal resistance at the 12-week-old stage, while Soraya showed intermediate lesion sizes compared to Durban and Rowena (Figure 1B-D).

IR was analysed after each elicitor application. A single treatment was insufficient to induce resistance in any of the cultivars (Supplementary Figure 1A). However, from the second application, resistance began to emerge in some treatments. In Durban, JA significantly reduced lesion size by 72.5% and 88.4% after the second and third applications, respectively (Supplementary Figure 1B, C). This effect was not sustained after the fourth application (Figure 1B), where none of the elicitors reduced lesion size in Durban. In contrast, in Rowena and Soraya, none of the elicitors induced statistically significant reductions in lesion size until the fourth application (Supplementary Figure 1A-C; Figure 1C, D). After the fourth application, whereas JA, SA, and I3CA did not induce resistance in Rowena, BABA treatment reduced lesion size by 41.6% compared to the Water control (Figure 1C). RBH was even more effective, reducing lesion size by 67.5% compared to Water (Figure 1C). In Soraya, RBH also significantly reduced lesion size by 64% after four applications (Figure 1D). However, none of the other elicitors, including BABA, result in a statistically significant difference in lesion size (Figure 1D).

To test the durability of these effects, additional assays were conducted on 18-week-old plants, six weeks after the final elicitor application. Differences in basal resistance between Rowena and Soraya at this developmental stage were not found (Figure 1E, F). In Rowena, BABA was the only elicitor to result in a reduced lesion size (45.3% compared to the control) (Figure 1E). In Soraya, RBH maintained a 42.6% reduction (Figure 1F). Interestingly, BABA and JA treatments also reduced lesion sizes by 44.8% and 40.7%, respectively (Figure 1F). These findings indicate that BABA, RBH, and JA can induce long-lasting resistance effects in specific cultivars.

### Relative Growth Rate analysis, flowering, and fruit production assessment

To evaluate whether repeated elicitor treatments impacted strawberry development, we assessed plant growth and reproductive traits following four consecutive applications from the seedling to the mature stage. No significant differences (*p* > 0.05) in RGR were observed across treatments in any of the three cultivars (Supplementary Figure 2). Flowering and fruit set were quantified by counting the number of flowers and fruits at 16 weeks of age (four weeks after the final elicitor application). No statistically significant effects (p > 0.05) of the elicitor treatments were found on flowering (Supplementary Table 1) or fruit (Supplementary Table 2) production in any cultivar.

### Metabolomic profiling of elicitor-induced resistance

To investigate the underlying mechanisms of induced resistance, an untargeted metabolomic analysis was conducted in Rowena and Soraya treated with Water, BABA, RBH, and JA, and then challenged with *B. cinerea* or a mock treatment. A total of 14,603 metabolomic features were detected, including 9,647 under positive electrospray ionization (ESI+) and 4,956 under negative mode (ESI−).

#### Variety-specific metabolomic profiles to infection

First, to investigate whether differences in basal resistance between cultivars could be associated with distinct constitutive metabolic profiles, we performed a PCA and hierarchical clustering heatmap analysis on mock- and *B. cinerea*-infected Water-treated plants (Figure 2). This was done to mirror the assessment shown in Figure 1C-D, where cultivar-specific differences in lesion size were observed. Figure 2A shows a PCA based on metabolomic features of Rowena and Soraya in the absence and presence of *B. cinerea*. The first two principal components (PC) explained 13.8% of the total variability, with the first component (PC1) accounting for 8,2% and the second component (PC2) accounting for 5.6%. While *B. cinerea* infection alone did not significantly alter the global metabolome at 24 hpi (PERMANOVA, F = 0.28, R² = 0.04, p = 0.942), orientation along PC1 reflects underlying metabolic differences between the two cultivars. The heatmap revealed a strong separation between Rowena and Soraya, with mock and infected samples clustered closely within each cultivar (Figure 2B). This supports the PCA results and confirms that infection had minimal impact on the global metabolic profile at 24 hpi, and that constitutive metabolomic differences exist between cultivars with contrasting basal resistance.

**Figure 2.**
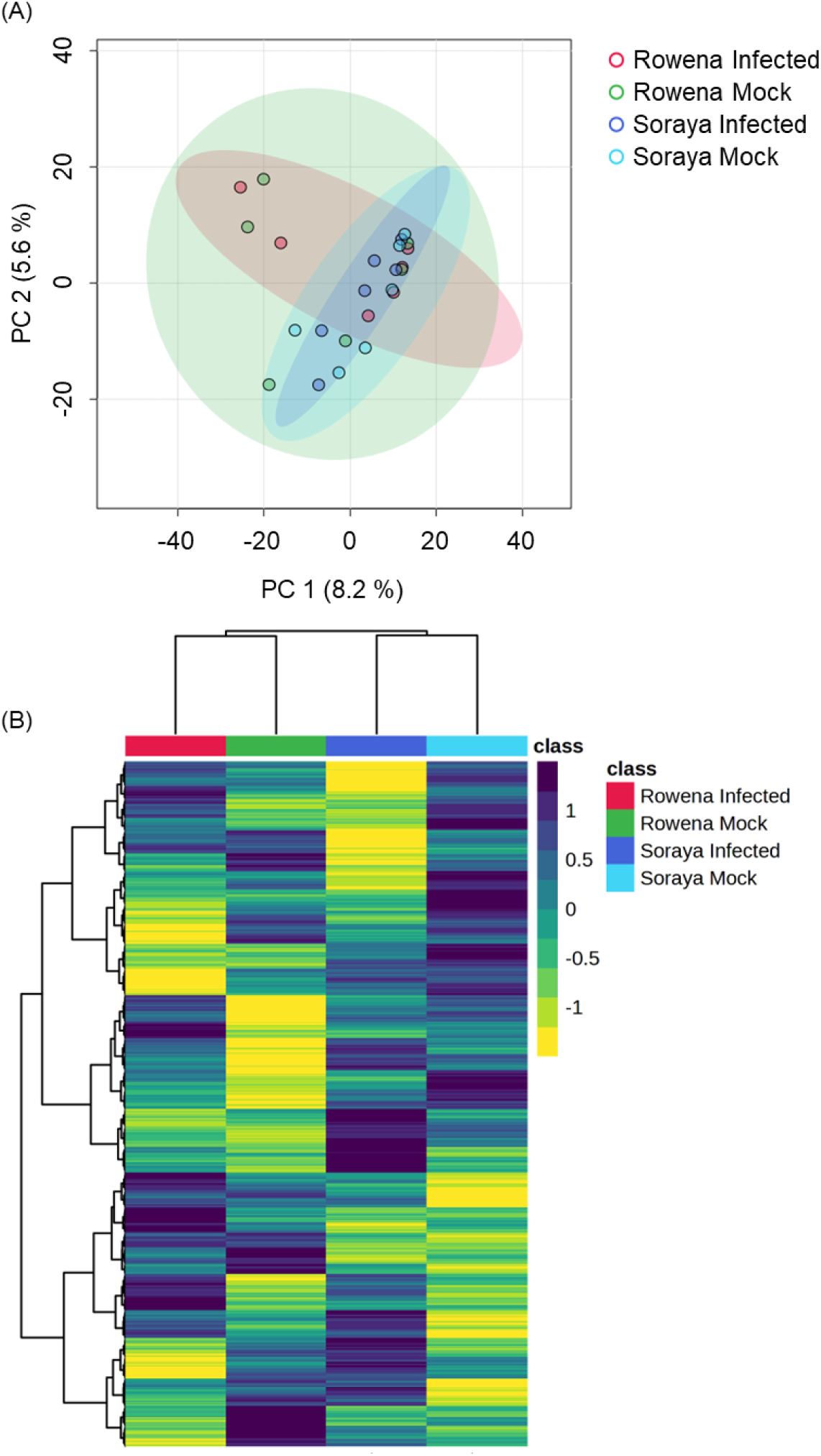
Basal metabolomic differences between strawberry cultivars with contrasting susceptibility to *B. cinerea*. (A) Principal Component Analysis (PCA) of metabolomic profiles in Water-treated Rowena and Soraya plants under mock- and *B. cinerea*-inoculated conditions. Samples were collected 24 hours post-inoculation. Each point in the PCA represents one replicate (n = six per condition; three biological replicates; two technical replicates). PERMANOVA was used to assess statistical significance. (B) Hierarchical clustering heatmap of metabolomic features from the same samples.

#### Metabolomic impact of BABA, RBH, and JA in Rowena and Soraya

Next, we assessed whether elicitor treatments triggered changes in the metabolic profiles of each cultivar and whether these changes were influenced by *B. cinerea* infection. In the PCA of Rowena samples under mock conditions, the first two PCs explained 24.5% of the total variability, with PC1 accounting for 13.6% and PC2 accounting for 10.9%. This PCA revealed clear separation between treatment groups (PERMANOVA, F = 5.66, R² = 0.45, p = 0.001; Figure 3A), with RBH showing the most distinct profile compared to Water, JA, and BABA. Under *B. cinerea* infection, PCs explained 28.9% of the total variability, with PC1 accounting for 17.9% and PC2 accounting for 11%. Again, it showed a significant treatment effect (PERMANOVA, F = 5.37, R² = 0.44, p = 0.001; Figure 3B). Notably, RBH remained the most metabolically distinct, while BABA showed a more evident divergence from Water upon infection than in the mock conditions.

**Figure 3.**
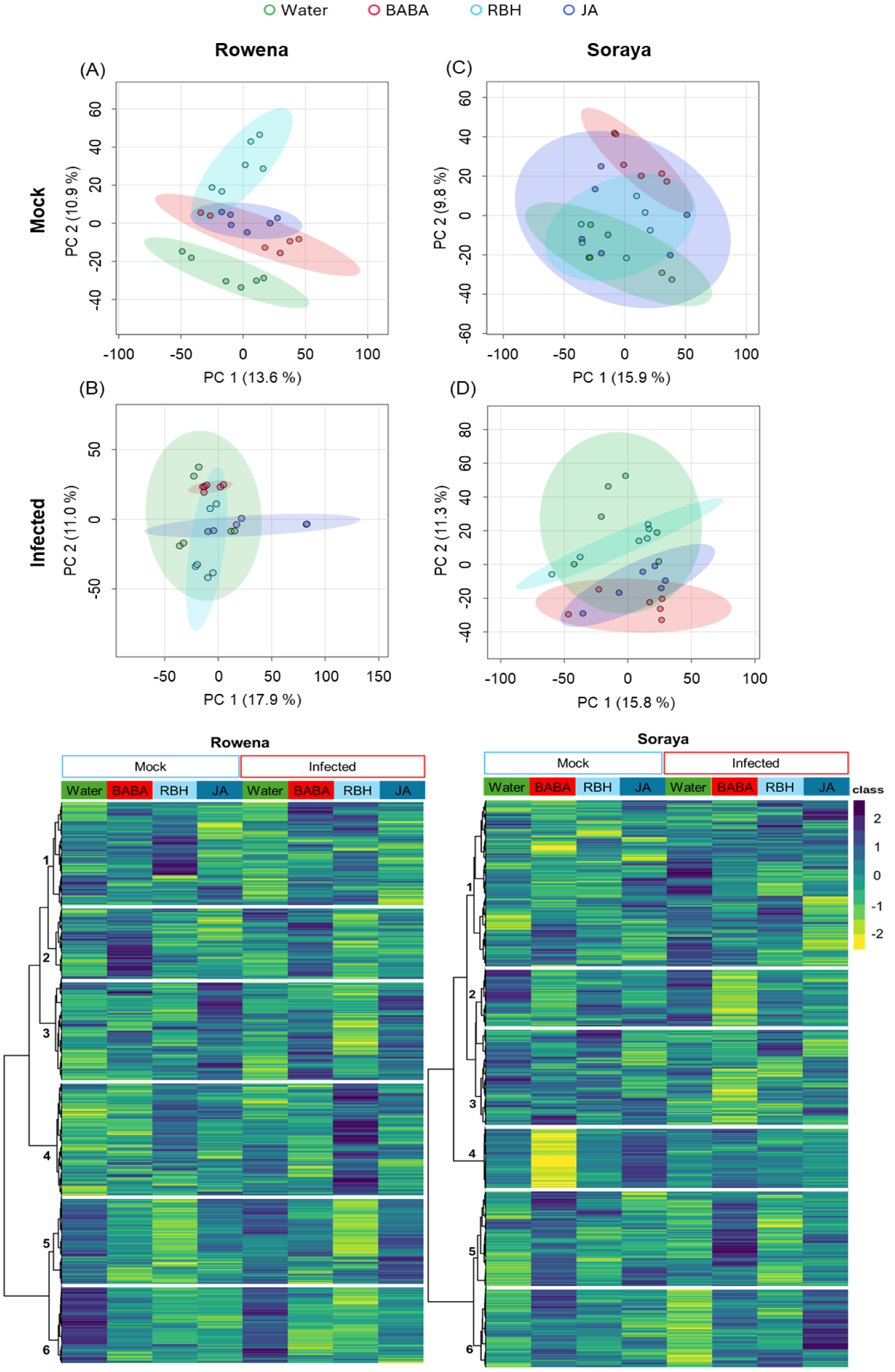
Metabolomic shifts in strawberry cultivars following elicitor treatments under mock and *B. cinerea*-inoculated conditions. Principal Component Analysis (PCA) of metabolic profiles in Rowena (A) and Soraya (C) under mock conditions and in Rowena (B) and Soraya (D) under *B. cinerea* infection. Samples were collected 24 hours post-inoculation. Each point represents one biological replicate (n = six per condition; three biological replicates; two technical replicates). Significant separation of treatment groups was observed by PERMANOVA (p < 0.05). (E) Hierarchical clustering heatmap of metabolomic features in Rowena across treatments (Water, BABA, RBH, and JA) under both mock and infected conditions. (F) Hierarchical clustering heatmap of metabolomic features in Soraya across the same treatments and conditions. Clusters highlight treatment- and cultivar-specific metabolic reprogramming.

In the PCA of Soraya samples under mock conditions, PCs explained 25.7% of the total variability, with PC1 accounting for 15.9% and PC2 accounting for 9.8%. Upon *B. cinerea* infection, PCs explained 27.1% of the total variability, with PC1 accounting for 15.8% and PC2 accounting for 11.3%. Similarly to the previous case, in Soraya, PCA also revealed significant treatment effects under both mock (PERMANOVA, F = 2.71, R² = 0.29, p = 0.027; Figure 3C) and infected conditions (PERMANOVA, F = 3.20, R² = 0.32, p = 0.012; Figure 3D). BABA had the most substantial impact, significantly altering the metabolome compared to Water and RBH in mock conditions and to Water under infection. These results suggest that elicitor-induced metabolic changes vary depending on both the cultivar and the presence of *B. cinerea*, with RBH being most effective in Rowena and BABA in Soraya.

A global view of elicitor-induced metabolic changes was further assessed through hierarchical clustering heatmap analysis, where the metabolic features detected in both ESI+ and ESI− modes were grouped according to treatment and compared between mock- and *B. cinerea*-inoculated conditions (Figure 3E-F).

In Rowena, clustering analysis revealed six significant metabolite clusters with expression patterns across treatments and infection conditions (Figure 3E). Cluster 1 included metabolites upregulated by RBH under mock conditions, downregulated by infection in Water-treated plants, and subsequently upregulated again in both BABA- and RBH-infected samples. Cluster 2 consisted of metabolites consistently upregulated by BABA under both mock and infected conditions. Cluster 3 showed metabolites upregulated by JA in both mock and infected samples, downregulated by RBH following infection (*i.e*., priming of RBH cluster), and an upregulated by BABA upon infection (*i.e*., priming of BABA cluster). Cluster 4 was defined by metabolites that were specifically upregulated after RBH treatment and infection (*i.e*., priming of the RBH cluster). Cluster 5 includes metabolites downregulated by RBH under both mock and infected conditions, with a greater impact of the infection (*i.e*., priming of RBH cluster). Cluster 6 represented metabolites that were consistently downregulated by all three elicitor treatments, regardless of the infection status.

In Soraya, six distinct clusters were identified (Figure 3F). Cluster 1 included metabolites with different profiles upon treatments and infection. Cluster 2 was associated with metabolites downregulated in BABA treatment and, to a much greater extent, in BABA-infected samples (*i.e.*, priming of the BABA cluster). Cluster 3 was defined by downregulation of metabolites after BABA treatment and infection (*i.e*., priming of the BABA cluster). Cluster 4 contained features downregulated in the BABA mock and upregulated in the JA mock. Cluster 5 captured metabolites that were upregulated under BABA mock conditions and to a much greater extent in BABA-infected samples (*i.e*., priming of the BABA cluster). Finally, Cluster 6 grouped features that were upregulated in BABA mock, downregulated in Water-infected plants, and upregulated in JA upon infection (*i.e*., priming of the JA cluster).

#### Isolation of direct and priming compounds and primed pathways identification in Rowena

To identify metabolites specifically associated with elicitor-induced resistance, we performed pairwise comparisons between each elicitor treatment and the Water control under both mock- and *B. cinerea*-inoculated conditions. Significant metabolites unique to either condition were categorised as “direct” (mock only) or “priming” (infection only).

In Rowena, when comparing all the putative metabolites under mock conditions, we observed that RBH triggered the most substantial metabolic reprogramming, with 348 (41.7%) induced metabolites, followed by JA (222, 26.6%) and BABA (32, 3.8%) (Supplementary Figure 3A). Shared metabolite changes included 64 between the three elicitors, 124 between RBH and JA, 27 between RBH and BABA, and 17 between JA and BABA. Upon *B. cinerea* infection, RBH continued to dominate the metabolic response, inducing 357 (58.8%) metabolites, followed by BABA (136, 22.4%) and JA (7, 1.2%) (Supplementary Figure 3B). Infected samples had no shared metabolites between the three elicitors, and only 107 were shared between BABA and RBH.

To isolate direct activators (elicitor-induced under mock conditions only), Venn diagrams between mock and infected for each elicitor showed 425 JA-, 339 RBH-, and 82 BABA-associated metabolites (Supplementary Figure 3C). These were further grouped to identify metabolites uniquely associated with each elicitor (Figure 4A). After this filtering (where the impact of the infection was removed), the effect of JA on the metabolome was more pronounced than the previously observed effect of RBH, with 306 (44.2%) metabolites being induced, followed by 221 (31.9%) RBH-induced and 32 (4.6%) BABA-induced (Figure 4A).

**Figure 4.**
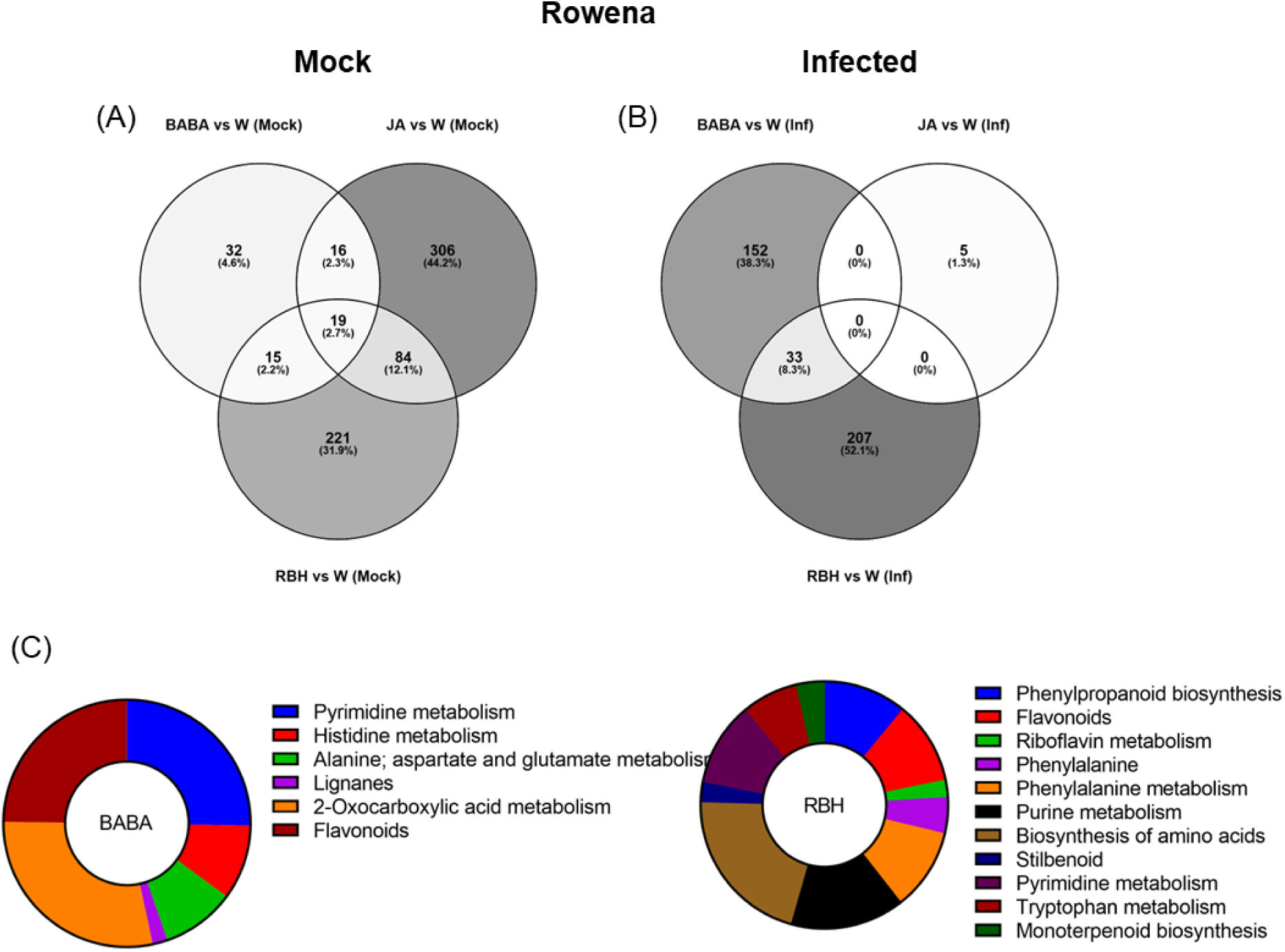
Metabolomic responses to elicitor treatments in Rowena. (A) Venn diagram showing the number of metabolites exclusively induced under mock conditions (no infection) by RBH, BABA, and JA in Rowena. (B) Venn diagram showing the number of metabolites exclusively induced under *B. cinerea* infected conditions by RBH, BABA, or JA in Rowena. Percentages represent each elicitor’s contribution to the total number of direct/priming-associated features. The number of shared metabolites between treatments is also indicated. (C) Pathway enrichment of priming-associated metabolites in Rowena. Putatively annotated priming-associated metabolites uniquely induced by BABA and RBH in *B. cinerea*-infected Rowena plants were subjected to pathway enrichment analysis using the KEGG database for *Fragaria vesca* and one internal library. Pie charts represent the percentage of statistically significant enriched pathways (p < 0.05) for BABA (left) and RBH (right), based on the number of annotated metabolites per pathway, are shown for each elicitor.

Priming-associated metabolites (elicitor-induced only upon *B. cinerea* infection) were also identified. Similar filtering was done to isolate metabolites associated only with the infection, which resulted in 240 RBH-specific, followed by 185 BABA- and 5 JA-associated features (Supplementary Figure 3C). After this filtering (where the impact of the direct effect of the elicitor was removed), we observed that 207 (52.1%) were exclusive to RBH, 152 (38.3%) to BABA, and 5 (1.3%) to JA. Only a shared group of 33 (8.3%) metabolites was found between BABA and RBH (Figure 4B).

Priming-associated metabolites from Rowena were selected based on their unique association with BABA, RBH, or JA treatments, as identified in the Venn diagrams. These metabolites were subjected to putative annotation by comparison with two reference libraries: the KEGG database for *Fragaria vesca* and an internal library kindly provided by Dr. Pastor’s group. Metabolite identifications were made in MassLynx using ChromaLynx, with confidence levels based on retention time and spectral matching (MS level). Where fragmentation spectra were not available in the selected libraries, complementary spectra were retrieved from public databases (e.g., PubChem, HMDB) using similar LC-MS/MS experimental parameters and ionization modes.

A total of 62 metabolites were putatively identified in Rowena: 23 were uniquely associated with BABA, 37 with RBH, and 2 were common to both treatments (Table 1). Enrichment analysis revealed statistically significant metabolic pathways related to these priming responses for BABA and RBH only. For BABA, enriched pathways included Alanine metabolism (3 metabolite; 2 up, 1 up), Flavonoids (5 metabolites, all down), Histidine metabolism (5 metabolites; 4 up, 1 down), Lignans (1 metabolite, up), Pyrimidine metabolism (4 metabolites; 3 up, 1 down), and 2-Oxocarboxylic acid metabolism (5 metabolites: 4 up, 1 down) (Figure 4c). RBH-associated priming metabolites were significantly enriched in Biosynthesis of amino acids (2 metabolites, down), Flavonoids (7 metabolites; 6 down, 1 up), Monoterpenoid biosynthesis (1 metabolite up), Phenylalanine-tyrosine-tryptophan biosynthesis (2 metabolites, 1 up, 1 down), Phenylalanine metabolism (4 metabolites; 3 down, 1 up), Phenylpropanoid biosynthesis (5 metabolites; 4 down, 1 up), Purine metabolism (7 metabolites; 4 down, 3 up), Pyrimidine metabolism (4 metabolites; 3 down, 1 up), Riboflavin metabolism (2 metabolites; 1 up, 1 down), Stilbenoid biosynthesis (3 metabolites; 2 up, 1 down), and Tryptophan metabolism (2 metabolites; 1 up, 1 down) (Figure 4C; Table 1).

**Table 1.**
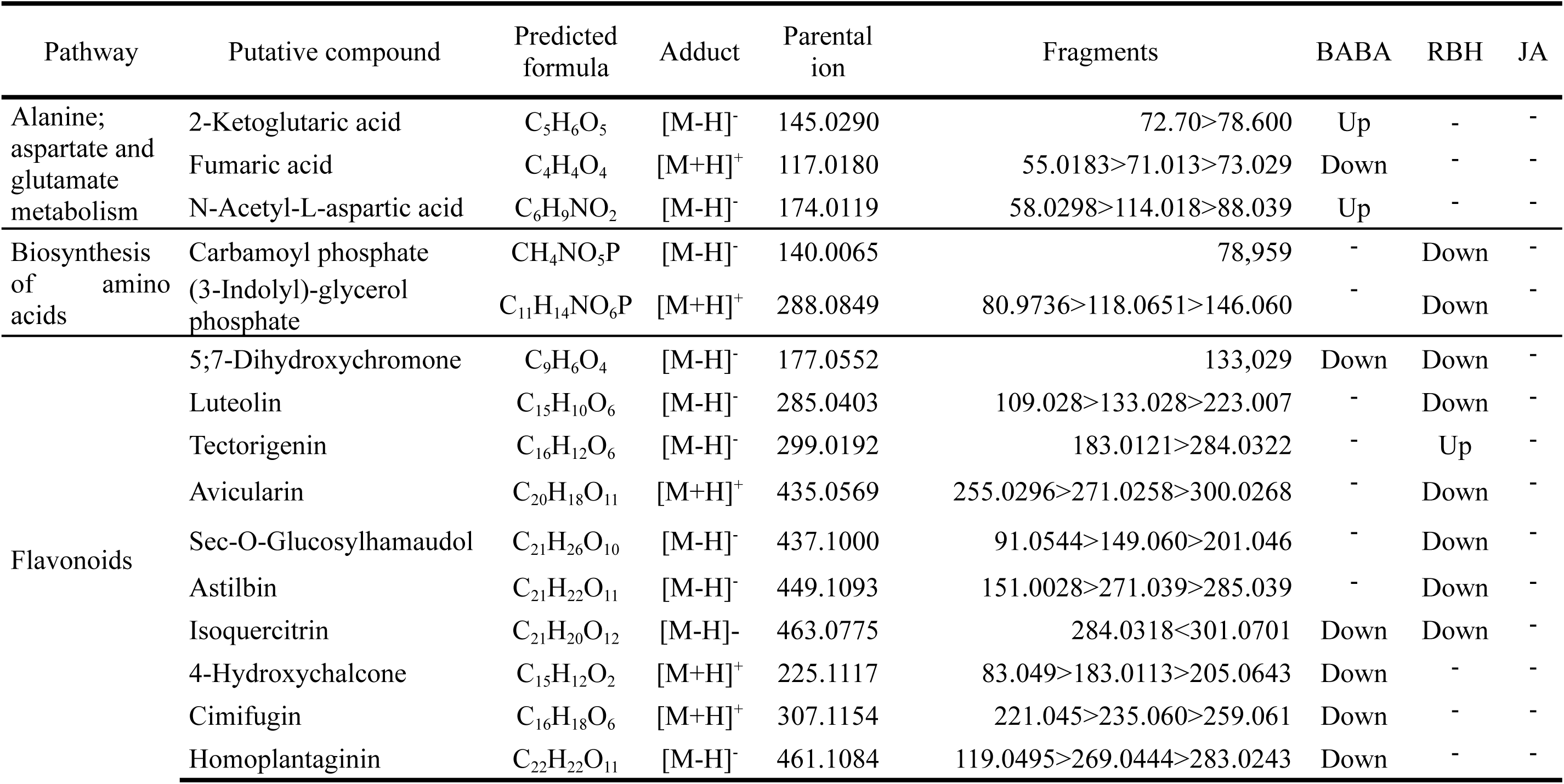
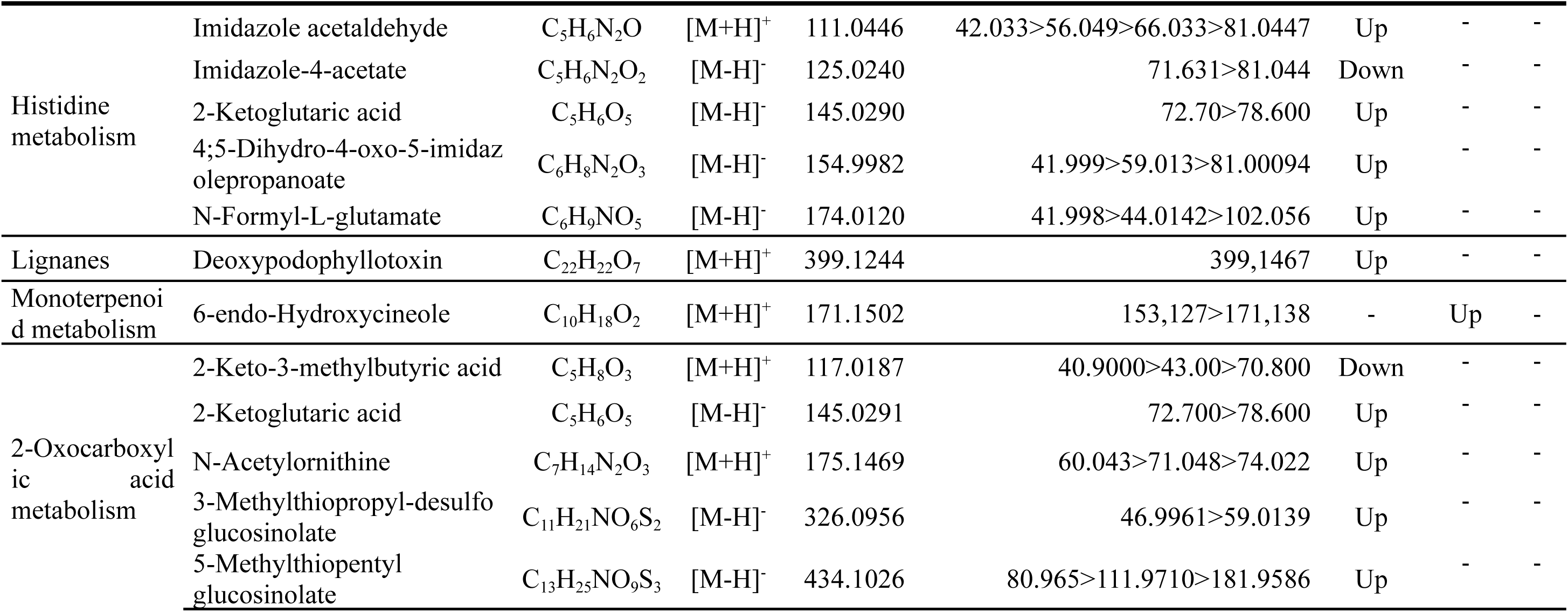
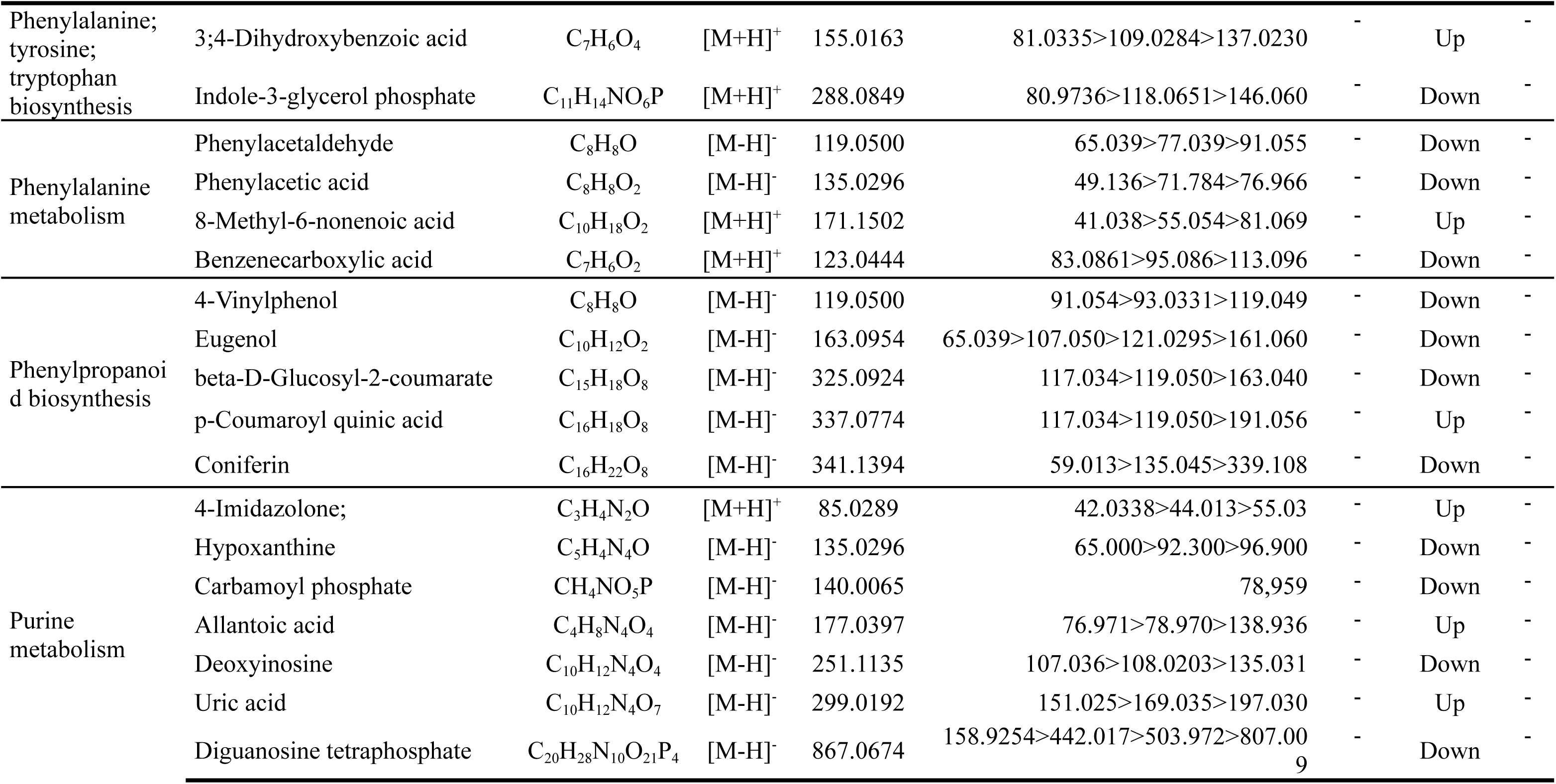
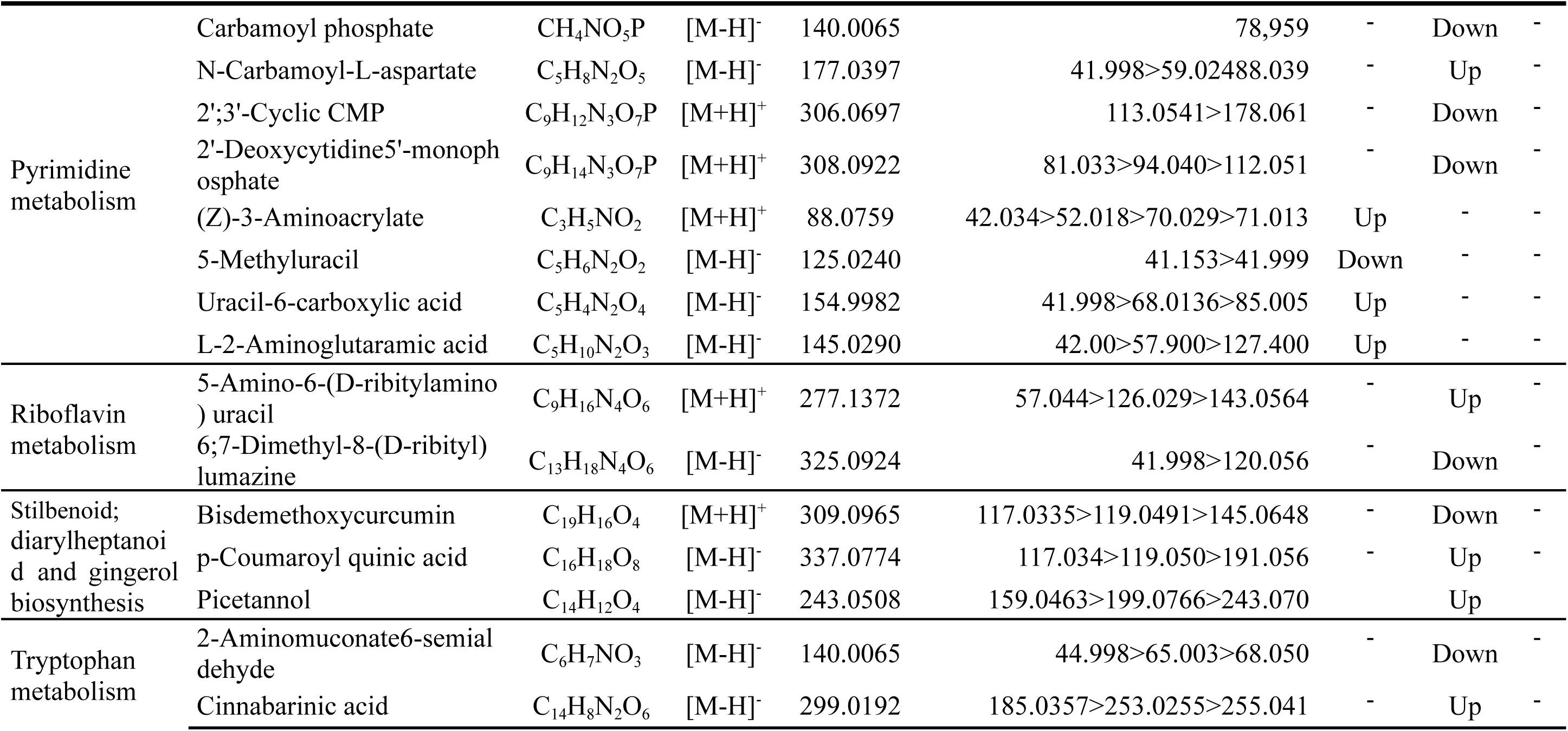
Summary of the pathways enriched by BABA, RBH and JA treatment in Rowena and the identified metabolites associated to those pathways. Up= metabolites upregulated; Down= metabolites downregulated compared to the corresponding treatment infected condition.

#### Isolation of direct and priming compounds and primed pathways identification in Soraya

A similar analysis was performed in Soraya. In contrast to Rowena, BABA triggered the most extensive metabolic reprogramming in Soraya under mock conditions, with 503 (97.3%) metabolites induced, whereas RBH and JA affected only 2 (0.4%) and 1 (0.2%) features, respectively (Supplementary Figure 4A). Minimal overlap was observed between treatments, with only 2 metabolites shared between BABA and RBH and 9 metabolites between BABA and JA. Upon *B. cinerea* infection, BABA continued to drive the strongest response, inducing 190 (41.1%) metabolites, followed by JA (103, 22.4%) and RBH (74, 16.1%) (Supplementary Figure 4B). Infected samples showed low shared responses, with 59 features shared between BABA and JA, 15 between BABA and RBH, 8 between JA and RBH, and only 10 features common to all treatments.

Direct-acting metabolites, *i.e*. those uniquely regulated under mock conditions, were predominantly BABA-associated (433), with JA and RBH contributing only 9 and 2 metabolites, respectively (Supplementary Figure 4C) This was also observed when comparing the filtered metabolites, where the dominance of BABA persisted with 425 (97.5%) metabolites remaining specific to this elicitor. In comparison, JA and RBH contributed 3 and 0 metabolites, respectively (Figure 5A).

**Figure 5.**
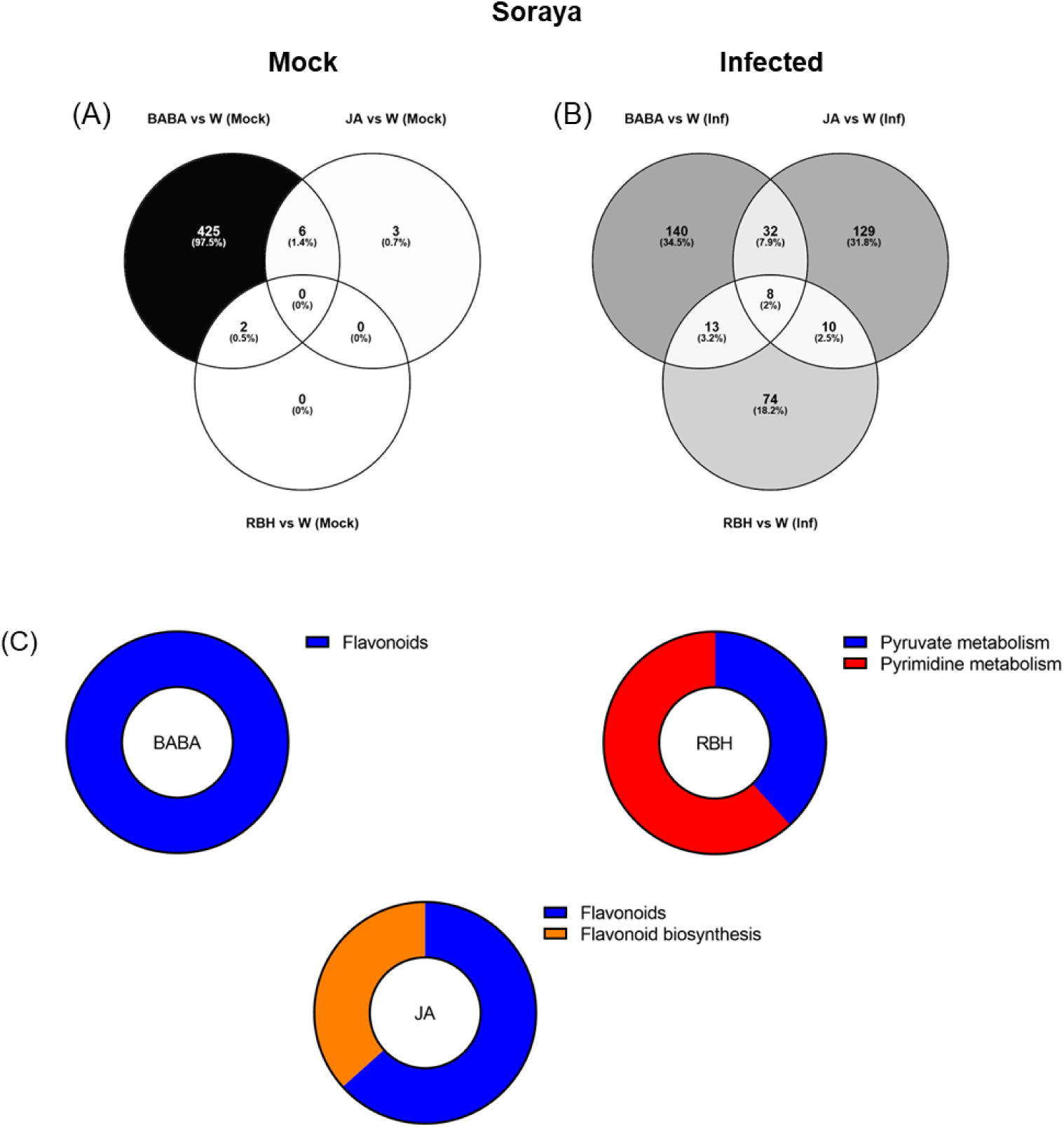
Metabolomic responses to elicitor treatments in Soraya. (A) Venn diagram showing the number of metabolites exclusively induced under mock conditions (no infection) by RBH, BABA, and JA in Soraya. (B) Venn diagram showing the number of metabolites exclusively induced under *B. cinerea* infected conditions by RBH, BABA, or JA in Rowena. Percentages represent each elicitor’s contribution to the total number of direct/priming-associated features. The number of shared metabolites between treatments is also indicated. (C) Pathway enrichment of priming-associated metabolites in Rowena. Putatively annotated priming-associated metabolites uniquely induced by BABA and RBH in *B. cinerea*-infected Soraya plants were subjected to pathway enrichment analysis using the KEGG database for *Fragaria vesca* and one internal library. Pie charts represent the percentage of statistically significant enriched pathways (p < 0.05) for BABA (left) and RBH (right), based on the number of annotated metabolites per pathway, are shown for each elicitor.

Priming-associated metabolites (elicitor-induced only in the presence of *B. cinerea*) revealed more balanced contributions. BABA was associated with 193 features, JA with 179, and RBH with 105 (Supplementary Figure 4C). Comparison of the priming effects by each elicitor revealed 140 metabolites (34.5%) exclusive to BABA, 129 (31.8%) to JA, and 74 (18.2%) to RBH (Figure 5B). The Venn diagram identified 8 metabolites (2%) shared by the three elicitors, 10 (2.5%) shared between JA and RBH, 32 (7.5%) shared between BABA and JA, and 13 (3.2%) shared between BABA and RBH. Pathway enrichment analysis was also performed in Soraya. A total of 16 priming-associated metabolites were putatively identified: 3 specific to BABA, 7 to RBH, and 5 to JA (Table 2). As with Rowena, metabolites were annotated using the KEGG database for *Fragaria vesca* and an internal reference library, and identifications were confirmed using ChromaLynx and MS spectral matching. Where in-library spectra were unavailable, additional spectral data were sourced from public databases such as PubChem and HMDB. Pathway enrichment analysis revealed that the Flavonoid biosynthesis pathway was significantly enriched for both BABA- and JA-associated priming metabolites, involving 3 metabolites (all upregulated) and 5 metabolites (3 upregulated, 2 downregulated), respectively. In contrast, RBH-primed metabolites were enriched in Pyruvate metabolism (4 metabolites all down) and Pyrimidine metabolism (3 metabolites all down) (Figure 5C; Table 2).

**Table 2.**
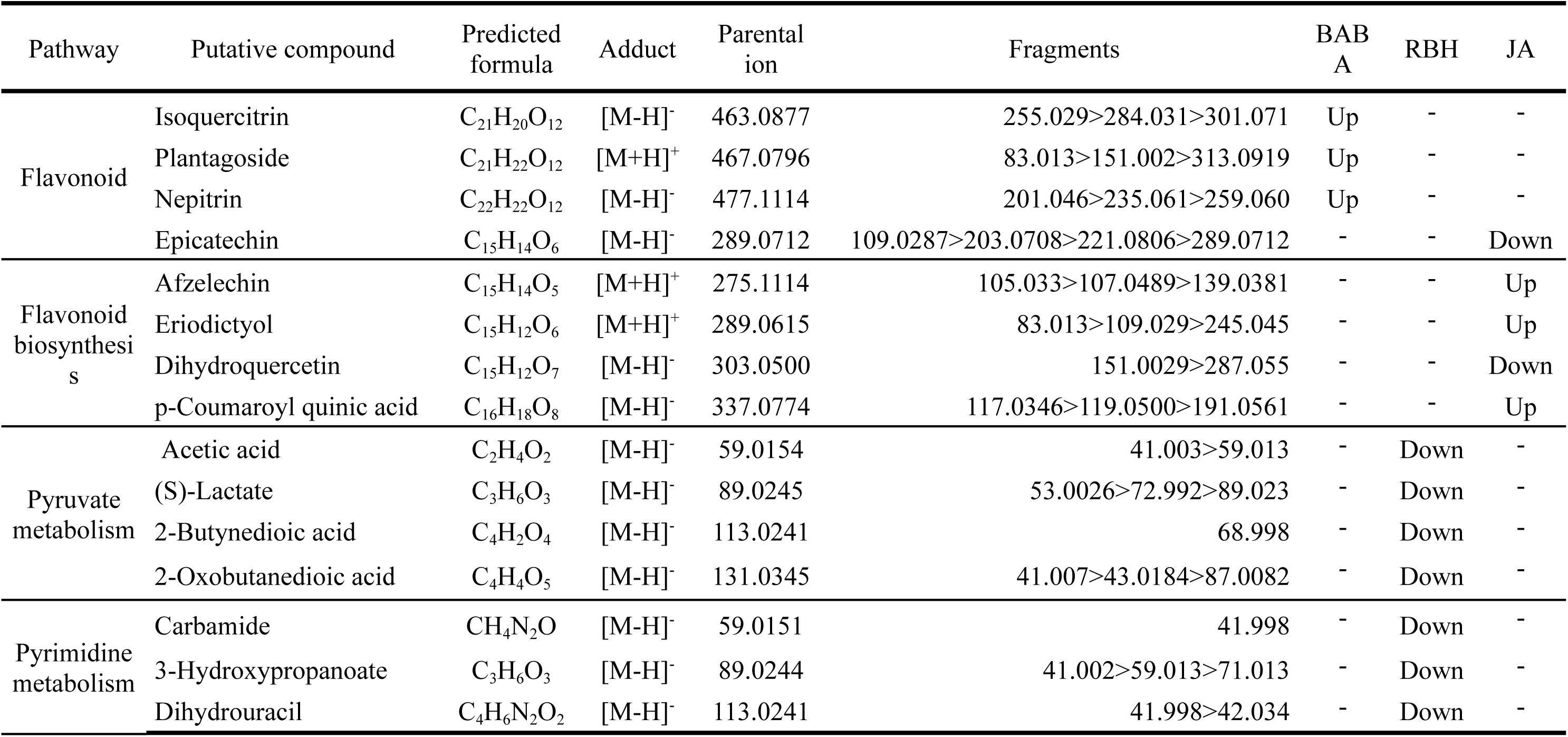
Summary of the pathways enriched by BABA, RBH and JA treatment in Soraya and the identified metabolites associated to those pathways. Up= metabolites upregulated; Down=metabolites downregulated compared to the corresponding treatment infected condition.

## Discussion

This study shows that the effectiveness of chemical elicitors in inducing resistance to *B. cinerea* in strawberry is both cultivar- and compound-dependent. Notably, RBH in Rowena and BABA in Soraya led to significant metabolome changes and reduced lesion development, even in long-term assessments. These findings highlight how cultivar-specific metabolic responses influence induced resistance, providing insight into potential pathways and biomarkers underlying the priming process.

Three commercial strawberry varieties, Durban, Rowena, and Soraya, were tested for their capacity to express induced resistance against *B. cinerea*. In strawberry, there are no genotypes fully resistant to *B. cinerea* (Bestfleisch et al., 2015, Petrasch et al., 2019). However, our results highlight differences in basal resistance among the three cultivars and across developmental stages. Durban and Rowena showed persistent susceptibility to *B. cinerea*, while Soraya exhibited greater resilience over time (Figure 1; Supplementary Figure 1). These differences, observed under controlled environmental conditions, suggest a genetic basis for basal resistance rather than an environmental influence.

We tested five chemical elicitors, BABA, RBH, I3CA, JA, and SA, for their capacity to induce resistance. Surprisingly, a single application of any elicitor was insufficient in all cultivars (Supplementary Figure 1A). This lack of induced resistance may be partially explained by the relatively low concentrations used in our study, which fall at the lower end of the ranges reported in successful induced resistance studies (Buswell et al., 2018). It is also likely that intrinsic, cultivar-specific traits modulate the responsiveness to these compounds. Strawberry cultivars exhibit significant variations in both basal and inducible defences, and previous studies have demonstrated differences in elicitor responsiveness among varieties (Seijo et al., 2008). Our findings reinforce the notion that genetic background plays a critical role in shaping immune responses in strawberries.

Neither I3CA nor SA conferred resistance in any of the cultivars. The ineffectiveness of I3CA could be hypothesized to be due to its potential to act more as a downstream marker of defence activation rather than as a true elicitor. Moreover, the lack of SA-induced resistance is not unexpected, as SA is typically associated with defence responses against biotrophic pathogens, whereas *B. cinerea* is a necrotroph. The antagonistic crosstalk between the SA and JA signaling pathways often results in SA being ineffective in necrotrophic pathogen resistance (Glazebrook, 2005, Spoel et al., 2007). Interestingly, earlier studies have shown that activation of the SA pathway can suppress JA-mediated responses, thus rendering plants more susceptible to *B. cinerea* (El Oirdi et al., 2011, Fugate et al., 2013, Ha et al., 2021, Khanam et al., 2005); however, we did not observe this in our experiments. We indeed observed some effects of JA, which significantly reduced lesion size in Durban after the second and third applications, consistent with its well-documented role in defence against necrotrophic pathogens.

The JA effect did not last after the third application and did not happen in Rowena or Soraya. This suggests that the concentration used may not have been strong enough to produce a strong response. Alternatively, this lack of effectiveness could be due to differences between the plant varieties in how they respond to hormones or how sensitive their receptors are to JA. These underlying factors highlight the complexity of elicitor interactions in strawberry and emphasize the importance of optimizing treatment conditions for each genetic background.

In contrast, RBH and BABA were highly effective in reducing lesion sizes in Rowena and Soraya after four applications, and their effectiveness continued beyond this point. RBH reduced lesion size by 67.5% in Rowena and 64% in Soraya, while BABA reduced lesion size by 41.6% in Rowena. Moreover, long-term resistance was assessed in 18-week-old plants. In Rowena, BABA continued to confer protection, while Soraya retained reduced lesion sizes with both BABA and RBH. This supports the capacity of these compounds to establish durable resistance, potentially through priming mechanisms, and these findings align with previous reports in tomato and *Fragaria vesca* (Badmi et al., 2019, Buswell et al., 2018, Wilkinson et al., 2017). However, BABA has been shown to induce susceptibility rather than resistance in *F. vesca*, with transcriptional profiling confirming this response (Badmi et al., 2022, Badmi et al., 2019). On the contrary, RBH has been demonstrated to induce resistance against *B. cinerea* in *F. vesca*, emphasising the differential activity of these compounds across Fragaria species (Badmi et al., 2019). Moreover, long-term induced resistance was also observed in treatments with JA in Soraya, which was unexpected due to the lack of short-term resistance induced by this elicitor in both Rowena and Soraya. This could be due to a delayed priming imprinting of the effects of the plant hormone, or to the developmental stage of Soraya. Durban showed no response to the elicitor treatments, likely due to genetic or physiological traits limiting its defence activation. Taken together, these results demonstrate the importance of species- and cultivar-specific responses to elicitors, suggesting that optimisation of elicitor use in strawberry will require consideration of both the compound and the cultivar.

Crucially, none of the elicitors impacted relative growth rate (RGR), flowering, or fruit production, suggesting a low risk of fitness penalties under our treatment conditions. This is particularly relevant for BABA, which has been associated with phytotoxicity and growth suppression in other species at higher doses (van Hulten et al., 2006). In contrast, RBH has shown minimal impact on growth and good systemic movement, supporting its potential use in sustainable agriculture.

Metabolomic analyses confirmed that the efficacy and mode of action of the elicitor are cultivar-dependent. First, we compared the metabolomic profiles of Rowena and Soraya after infection. PCA and hierarchical clustering of Water-treated plants revealed clear separation between Rowena and Soraya, independent of infection status (Figure 2). At 24 hpi, *B. cinerea* had little effect on the global metabolome, demonstrating that cultivar identity was the dominant factor influencing baseline metabolic variation. These metabolomic distinctions may underlie the differential responses to elicitor-induced resistance observed in our study.

Next, we examine the impact of specific elicitors on the metabolome of both varieties (Figure 2). In Rowena, RBH had the most pronounced metabolic impact under both mock and infected conditions, indicating a direct and priming effect. In Rowena, the BABA effect was infection-dependent, suggesting a priming-specific response. In Soraya, however, BABA had strong effects under both conditions, consistent with a mechanism of direct activation. JA exhibited minimal influence on Rowena but did alter the Soraya metabolome during infection, indicating an infection-dependent mode of action.

Venn diagrams and PCA-supported heatmaps revealed that RBH in Rowena and BABA in Soraya triggered the most substantial metabolic shifts (Figure 4-5). In both cases, these elicitors led to a higher diversity and abundance of priming-associated metabolites, which correlated with increased resistance phenotypes. This indicates that elicitor efficacy depends on the plant’s ability to adjust its metabolism in response to both treatment and pathogen. In Soraya, while the four applications of BABA did not significantly reduce lesion size, there was a noticeable trend towards improved resistance. Notably, long-term resistance was strongly induced following BABA treatment, suggesting that while the direct activation of defence responses may be transient or subtle, durable priming effects are maintained over time. This supports a model in which BABA acts through sustained priming rather than immediate defence activation in Soraya.

Pathway enrichment analysis provided key insights into elicitor-induced resistance mechanisms (Figure 4-5). In Rowena, RBH led to significant enrichment in various primary metabolic pathways, including amino acids, nucleotides, cofactors, and vitamins.

Amino acid metabolism was significantly impacted in Rowena, especially in the phenylalanine, tyrosine, and tryptophan pathways, with most metabolites generally downregulated upon RBH treatment. Similarly, enrichment of purine and pyrimidine metabolism also occurred, with most metabolites downregulated. This downregulation of primary metabolism under RBH treatment may suggest reallocations of energy resources during priming, redirected towards defence pathways (Rojas et al., 2014). Accordingly to our results, amino acid metabolism was found to be suppressed during priming (Schwachtje et al., 2019). Cofactors and vitamins metabolism enriched by RBH in Rowena, included precursors of riboflavin (Vitamin B2), whose efficacy as elicitor of systemic resistance against fungal pathogens (including *B. cinerea*) relies on activation of Reactive Oxygen Species (ROS) signalling, activation of lipoxygenase pathway, pathogenesis-related proteins (PR), and defence and antioxidant enzymes such as phenylalanine ammonia lyase (PAL) and peroxidase (POD) (Azami-Sardooei et al., 2010, Boubakri et al., 2013, Zhu et al., 2024). Within secondary metabolism, monoterpenoid and stilbenoid biosynthesis were upregulated by RBH in Rowena. Monoterpenoids can kill the pathogen directly or, due to their volatile properties, act as long-distance signals to trigger defence responses in the distal parts of the plant (Riedlmeier et al., 2017). Stilbenoids’ role against *B. cinerea* has been widely studied, especially for resveratol (Ahn et al., 2015, Xu et al., 2018). In our study, Piceatannol, a hydroxylated derivative of resveratrol, was found to be upregulated. Surprisingly, flavonoid biosynthesis was largely downregulated in Rowena under RBH treatment. Although most of the metabolites in phenylpropanoid-linked pathways were downregulated, p-Coumaroyl quinic acid was the only one upregulated. This metabolite is not only involved as an intermediate in flavonoid and lignin pathways, but also plays a central role in the antioxidant and ROS-scavenging mechanisms (Lee et al., 2013).

Unlike for RBH, BABA-primed Rowena plants showed a comparable enrichment of primary metabolism, particularly with upregulation in amino acid (e.g., histidine, alanine, aspartate, glutamate) and nucleotide (e.g., pyrimidine) metabolism, as well as 2-oxocarboxylic acid metabolism. Our results support previous findings by Pastor et al. (2014), which demonstrate that the influence of BABA on primary metabolism during the priming phase includes the accumulation of amino acids and enhanced Tricarboxylic acid (TCA) cycle activity. Similarly to Pastor et al. (2014), TCA intermediates, 2-oxoglutaric acid and Fumarate, were enriched under BABA treatment. Interestingly, both pyrimidine biosynthesis and degradation pathways were upregulated within nucleotide metabolism, suggesting the maintenance of nucleotide homeostasis. This is notable given that the BABA receptor in *Arabidopsis* has been identified as an aspartyl-tRNA synthetase, linking BABA perception to amino acid metabolism and protein synthesis (Luna et al., 2014a). Allantoic acid, uric acid, and 4-imidazolone, within catabolism of purine for storage and transport of nitrogen, were upregulated, suggesting involvement of BABA in Nitrogen metabolism (Ohyama et al., 2023, Yang and Han, 2004).

Similarly to the effect of RBH, flavonoid and lignan biosynthesis were also enriched, with most flavonoids downregulated and lignans upregulated. During the priming phase, plants can accumulate and store conjugates of compounds that play a role in defences (Pastor et al., 2014). According to this, we found that BABA and RBH induced the enrichment of the glucoside form of quercetin, Isoquercitrin, which is hydrolysed by plant β-glucosidases, releasing quercetin, a potent antioxidant, and scavenging reactive oxygen species (ROS), thereby activating plant immune responses. For instance, pre-treatment with quercetin induced resistance through the SA-dependent signalling pathway in *Arabidopsis* against *Pseudomonas syringae,* leading to the conclusion that similar flavonoids (similar to quercetin) may also work (An et al., 2023). These results highlight common metabolic signatures for BABA and RBH in Rowena, consistent with an earlier study (Buswell et al., 2018) and reinforce the idea that both compounds activate similar metabolic pathways to improve resistance in this cultivar.

In conclusion, as a priming agent, RBH in Rowena suppresses primary metabolism at the early stage of the infection and induces selective secondary metabolism (upregulation of stilbenoids and monoterpenoids). At the same time, BABA has a broader metabolic impact, especially on primary metabolism, which is upregulated and induces selective secondary metabolism (upregulation of lignanes). These results highlight different fine-tuning metabolic reprogramming performed by RBH and BABA in Rowena.

In Soraya, pathway enrichment analyses revealed distinct metabolomic signatures for each elicitor, with BABA exerting a powerful impact. BABA-primed metabolites were mainly enriched in flavonoid biosynthesis. Specifically, several flavonol glycosides, including Isoquercitrin, Plantagoside, and Nepitrin, were upregulated. Upregulation of Isoquercitrin in BABA-treated Rowena and RBH-treated Soraya suggests a potential role as a conserved marker of elicitor responsiveness. JA-priming in Soraya also affected flavonoid metabolism. While some precursors, such as Eriodictyol and compounds like Afzelechin and *p*-Coumaroyl quinic acid, were upregulated, key flavonoids such as Epicatechin and its precursor Dihydroquercetin were downregulated, indicating a complex regulation of this pathway during JA-induced priming. RBH treatment in Soraya induced enrichment of several secondary metabolic pathways, including phenylpropanoid, flavonoid, and stilbenoid biosynthesis. Within these, 5,7-Dihydroxychromone and Isoquercitrin were consistently modulated.

In contrast to Rowena, RBH-primed Soraya plants showed downregulation of primary metabolic pathways, including nucleotide (pyrimidine) and carbohydrate (pyruvate) metabolism. The downregulation of pyruvate metabolism may reflect energy conservation strategies associated with priming. Similarly, suppression of pyrimidine degradation pathways, which are crucial for nucleotide recycling and homeostasis, may represent a stress-mediated reprogramming strategy that contributes to effective resistance. Overall, these findings highlight that BABA and RBH induce distinct but partially overlapping metabolic responses in Soraya, with flavonoid biosynthesis emerging as a common target. The differential modulation of energy-related and defence-associated pathways illustrates how elicitor-induced priming operates through cultivar-specific metabolic adaptations.

Together, our results show that BABA, RBH, and JA induce distinct metabolomic reprogramming depending on the cultivar. While RBH suppresses primary metabolism but activates different secondary pathways in both cultivars, BABA elicits primary metabolism in Rowena and secondary metabolism in Soraya. The JA effect is weak and absent in Soraya but present in Rowena. The lack of observed growth penalties further highlights the potential of these elicitors in sustainable disease management strategies for strawberry. This study provides a comprehensive metabolomic view of defence priming induced by BABA, RBH, and JA in commercial strawberry cultivars. By integrating phenotypic and metabolomic data, we reveal how cultivar-specific responses and metabolic reprogramming influence the efficacy of elicitors. These insights can inform the strategic deployment of chemical elicitors in strawberry breeding and disease management programs to combat *B. cinerea* in a sustainable way.

## Author contribution

CM conducted all experiments, analysed the data, and wrote the manuscript. VP supervised the metabolomic analyses, provided technical support during the metabolomics experiments, guided CM in metabolomic data interpretation and contributed to metabolite identification. TRF co-supervised CM, secured project funding and offered intellectual input throughout the study. SMPC, as CM’s primary supervisor, provided continuous mentorship, secured project funding, contributed to the study’s conceptual development and offered critical intellectual guidance. EL co-supervised CM, contributed to the design and execution of experiments, supported experimental logistics and funding, co-wrote the manuscript and provided key intellectual input at all stages of the project.

## Acknowledgements

The authors would like to thank the Foundation for Science and Technology (FCT) for funding CM Ph.D. scholarship (2021.06641.BD) and the R&D Project ‘Botrytis-XTalk’ (PTDC/ASP-PLA/4478/2021) to TRF and SMPC. This research was also supported by national funds via FCT through the Strategic Projects UID/05748/2020. We also thank the Servicio de Instrumentación Científica (SCIC) at UJI for its technical support and the project CI AICO/2021/092 from the Generalitat Valenciana. The work was also supported by the BBSRC Future Leader Fellowship (BB/P00556X/2) and the pump-priming funding received by the Horticultural Quality and Food Loss Network (WXA3189N/P16188/UoB_Luna-Diez) to EL. We also thank Lamya Majeed for the technical experimental support to CM and to Saturn Bioponics for providing the commercial strawberry cultivars.

## Supplementary Figures

**Supplementary Figure 1.**
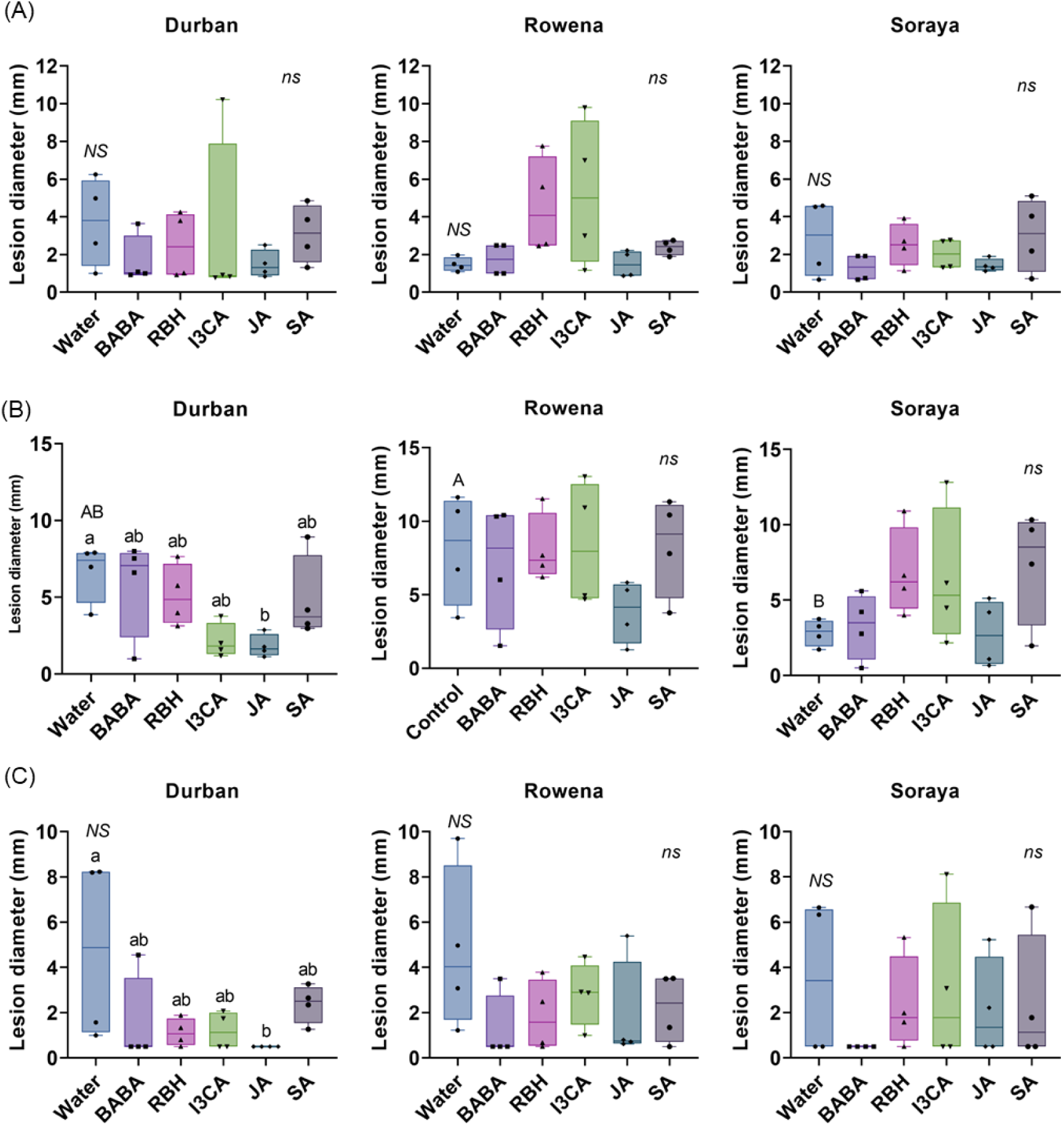
Lesion development in strawberry cultivars during early elicitor applications. Lesion diameter in Durban, Rowena, and Soraya after the first (A), second (B), and third (C) elicitor applications at 6, 8, and 10 weeks of plant age, respectively. Boxplots represent lesion diameter (mm) with the median line, interquartile range (boxes), and whiskers extending to the minimum and maximum values. Each point represents biological replicates (individual plants) (n = 8–12 per treatment). Capital letters indicate statistical differences between the Water treatments of each variety. Lowercase letters indicate statistically significant differences between elicitors within each variety (One-way ANOVA followed by Tukeýs post hoc test or Welch’s ANOVA followed by Dunnett’s T3; p < 0.05; n = 8–12).

**Supplementary Figure 2.**
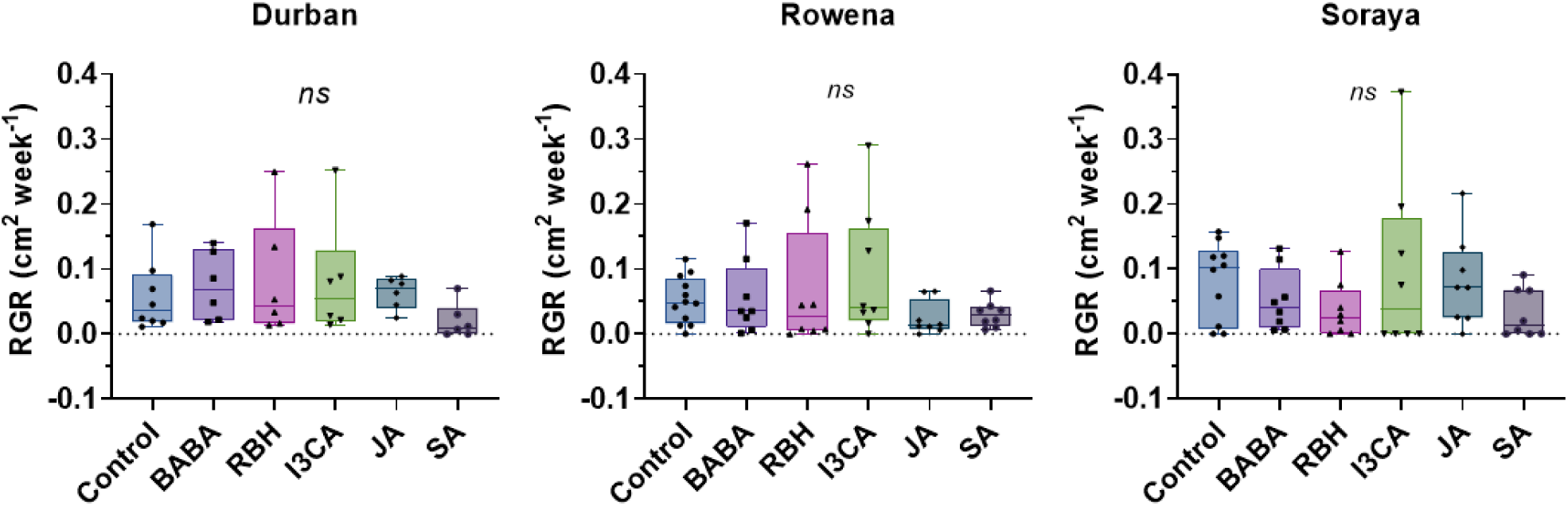
Assessment of growth following elicitor treatments. Relative Growth Rate (RGR) was calculated based on leaf area expansion between weeks 11 and 14 using ImageJ software. Boxplots represent RGR (cm^2^ week^-1^) with the median line, interquartile range (boxes), and whiskers extending to the minimum and maximum values. Each point represents biological replicates (individual plants). *ns* indicates not significant differences (One-way ANOVA; p < 0.05; n = 8–12).

**Supplementary Table 1.** Number of flowers counted for each treatment in each cultivar, after the third elicitor application (week 11 of the experiment) and the fourth elicitor application (week 13 of the experiment).

| Time<br>(week) | Cultivar | Water | BABA | RBH | I3CA | JA | SA |
| --- | --- | --- | --- | --- | --- | --- | --- |
| 11 | Durban | 33 | 25 | 10 | 16 | 22 | 19 |
|  | Rowena | 13 | 10 | 8 | 17 | 6 | 11 |
|  | Soraya | 24 | 9 | 24 | 16 | 22 | 9 |
| 13 | Durban | 26 | 24 | 13 | 19 | 20 | 17 |
|  | Rowena | 8 | 11 | 9 | 13 | 10 | 8 |
|  | Soraya | 23 | 10 | 18 | 14 | 21 | 7 |

**Supplementary Table 2.** Number of fruits counted for each treatment in each cultivar, after the fourth elicitor application (week 16 of the experiment). Nd: not detected

| Cultivar | Water | BABA | RBH | I3CA | JA | SA |
| --- | --- | --- | --- | --- | --- | --- |
| Durban | 7 | nd | nd | 1 | 4 | 4 |
| Rowena | 6 | 4 | nd | 4 | 1 | 7 |
| Soraya | 9 | nd | nd | 7 | 6 | 5 |

**Supplementary Figure 3.**
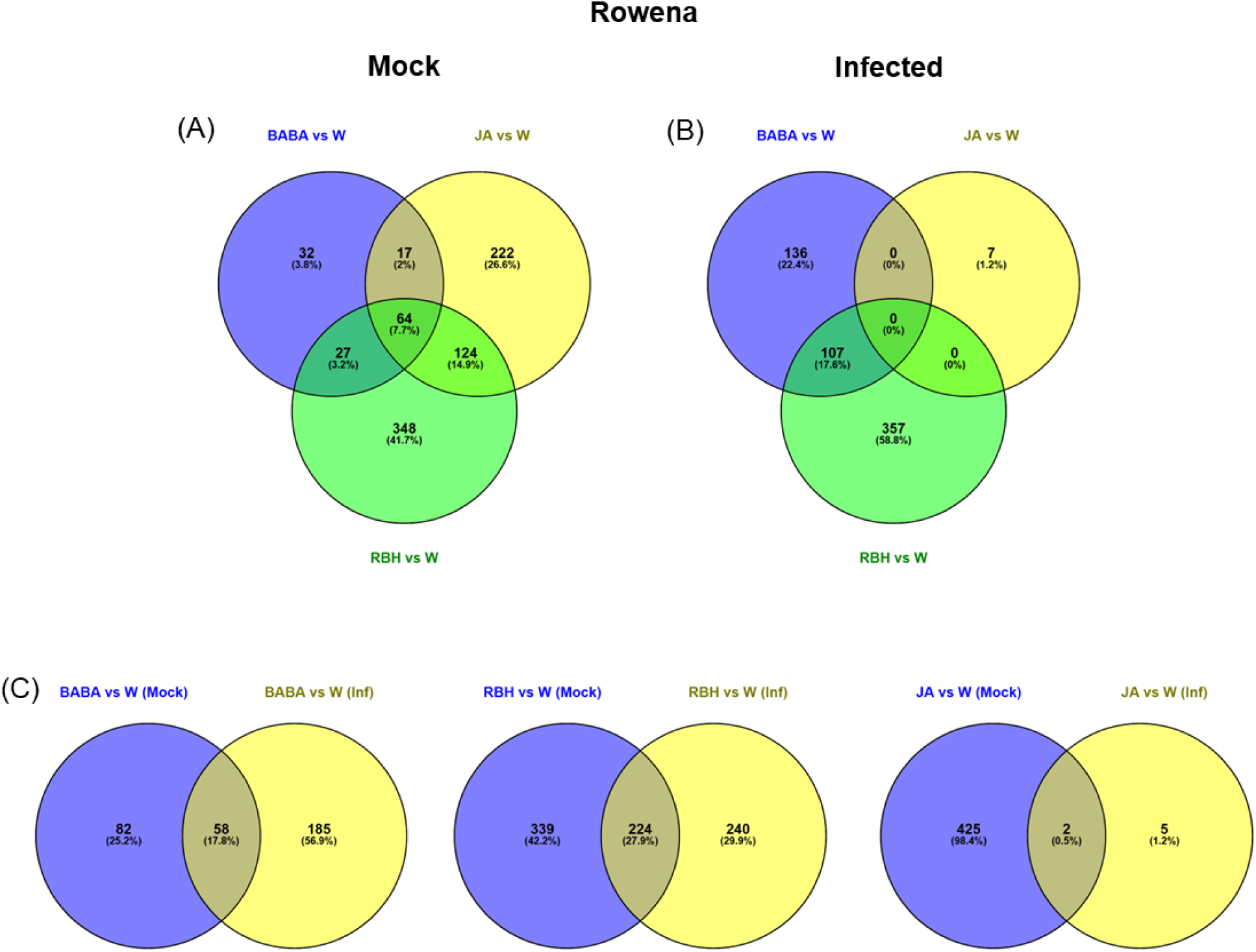
Metabolite reprogramming in Rowena in response to elicitors. (A) Venn diagram showing metabolites induced under mock condition by BABA, RBH, and JA in Rowena. (B) Venn diagram showing metabolites induced under *B. cinerea* infected condition. (C) Filtering of mock-(blue circles) and infection (yellow circles)-specific metabolites for each treatment. The diagram separates direct (mock-only) and priming (infection-only) responses, prior to exclusive grouping in Figure 4.

**Supplementary Figure 4.**
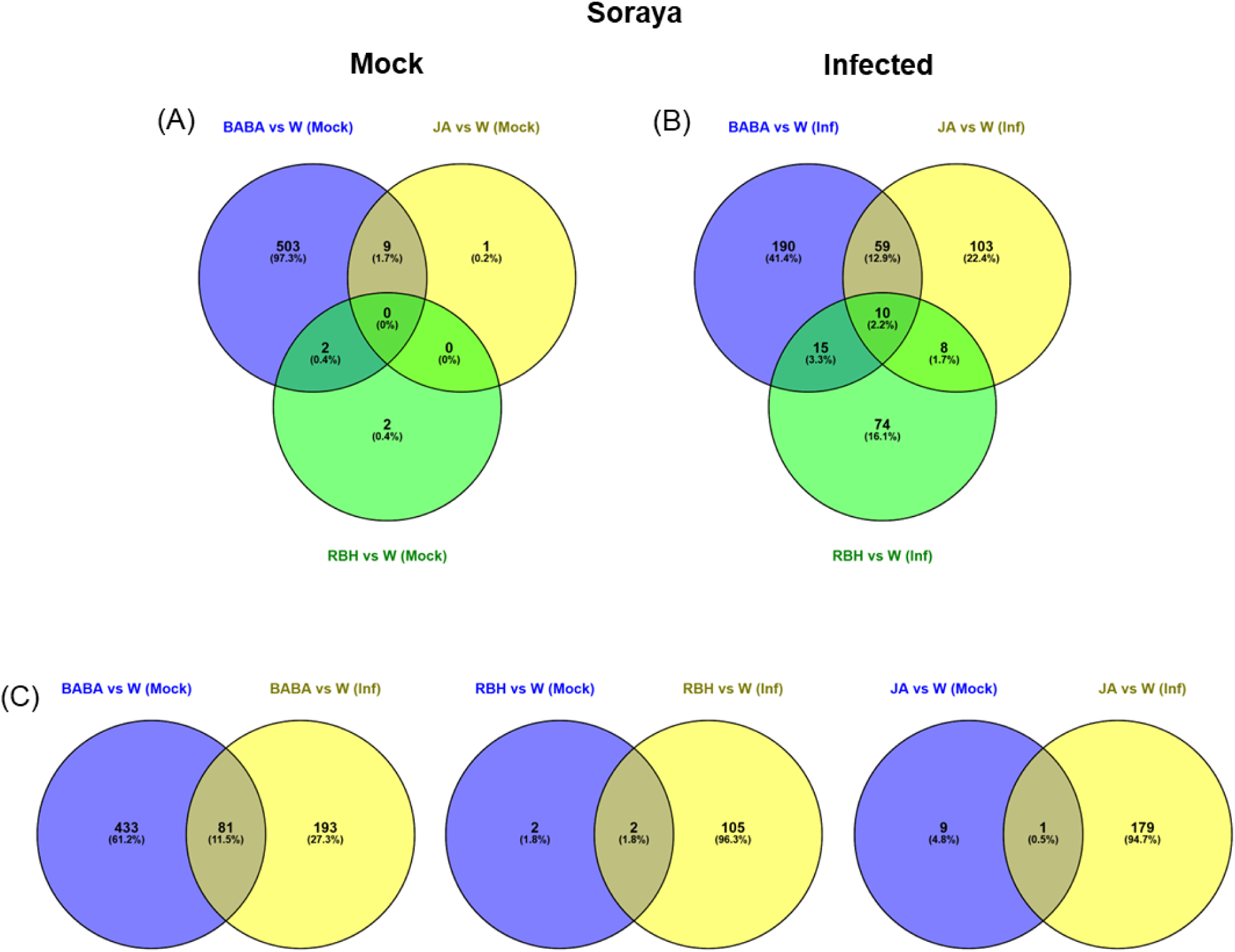
Metabolite reprogramming in Soraya in response to elicitors. (A) Venn diagram showing metabolites induced under mock condition by BABA, RBH, and JA in Soraya. (B) Venn diagram showing metabolites induced under *B. cinerea* infected condition. (C) Filtering of mock-(blue circles) and infection (yellow circles)-specific metabolites for each treatment. The diagram separates direct (mock-only) and priming (infection-only) responses, prior to exclusive grouping in Figure 5.

